# Mediator Subunit Med4 Enforces Metastatic Dormancy in Breast Cancer

**DOI:** 10.1101/2023.11.18.566087

**Authors:** Seong-Yeon Bae, Hsiang-Hsi Ling, Yi Chen, Hong Chen, Dhiraj Kumar, Jiankang Zhang, Aaron D. Viny, Ronald A. DePinho, Filippo G. Giancotti

## Abstract

Long term survival of breast cancer patients is limited due to recurrence from metastatic dormant cancer cells. However, the mechanisms by which these dormant breast cancer cells survive and awaken remain poorly understood. Our unbiased genome-scale genetic screen in mice identified *Med4* as a novel cancer-cell intrinsic gatekeeper in metastatic reactivation. *MED4* haploinsufficiency is prevalent in metastatic breast cancer patients and correlates with poorer prognosis. Syngeneic xenograft models revealed that *Med4* enforces breast cancer dormancy. Contrary to the canonical function of the Mediator complex in activating gene expression, *Med4* maintains 3D chromatin compaction and enhancer landscape, by preventing enhancer priming or activation through the suppression of H3K4me1 deposition. *Med4* haploinsufficiency disrupts enhancer poise and reprograms the enhancer dynamics to facilitate extracellular matrix (ECM) gene expression and integrin-mediated mechano-transduction, driving metastatic growth. Our findings establish *Med4* as a key regulator of cellular dormancy and a potential biomarker for high-risk metastatic relapse.

## Introduction

Survivorship for patients with breast cancer is predicated not by the disease burden in the breast itself, but rather the ability of cancer cells to metastasize outside the breast. Clinical experience and experimental data in both patients and mouse models show that metastatic disease is the overwhelming cause of breast cancer deaths. Cellular and molecular studies of metastatic breast cancer suggest that disseminating cells from the primary tumor undergo a variable period of dormancy at pre-metastatic sites before undergoing reactivation and outgrowth becoming macroscopic and life-threatening lesions(Early Breast Cancer Trialists’ Collaborative, 2005). These data point to a distinct stage of tumor progression whereby quiescence may serve as both a biomarker of high-risk disease as well as a key therapeutic target. We posit that cancer cell dormancy within the early metastatic niche provides a therapeutic window of opportunity across breast cancer subtypes including triple-negative breast cancer (TNBC), despite its historically perceived short clinical latency(Giancotti, 2013; Zhang et al., 2013),with a subgroup of TNBC patients exhibiting recurrence after five years(Rueda et al., 2019).

Genomic studies of breast cancer growth kinetics have indicated that systemic spread can occur rapidly following malignant transformation(Hosseini et al., 2016; Hu et al., 2020; Turajlic and Swanton, 2016; Werner-Klein et al., 2020). Early disseminated cancer cells (DCCs) often lack critical genetic and genomic alterations, which they need to acquire at distant sites. This suggests that their outgrowth in target organs may be driven by epigenetic mechanisms regulating key features of cellular identity, such as epithelial-to-mesenchymal transition (EMT) and stemness/self-renewal(Lambert et al., 2017). Indeed, analysis of metastasis driver genes through gene ontology has revealed enrichment for chromatin regulators in metastatic progression(Hu et al., 2020). Despite their clinical importance, metastatic dormancy and reactivation are poorly understood due to the complexity of the underlying biology. To untangle this complexity, we developed a forward genetic screening strategy in the mouse to uncover key regulators of metastatic reactivation(Gao et al., 2014). Screening of open reading frame (ORF) libraries enabled us to identify several positive regulators of metastatic reactivation of breast cancer, including Coco, which binds to and inhibits BMP in the lung microenvironment, and the tetraspanin family member TM4SF1, which mediates non-canonical activation of the JAK2-STAT3 signaling pathway(Gao et al., 2012; Gao et al., 2016).

Collectively, our prior studies suggest that metastatic outgrowth is initiated by a subpopulation of cancer stem cells with metastatic potential (metastasis-initiating cells), which undergo dormancy and reactivation in response to then activation of signaling pathways and transcriptional networks similar to those that govern these processes in adult stem cells(Giancotti, 2013). However, numerous modulators involved in regulating metastatic dormancy remain to be identified due to the extensive heterogeneity observed in the molecular, genetic, and clinical characteristics of breast cancer metastases. To identify additional modulators associated with metastatic dormancy, we conducted an *in vivo* CRISPR-Cas9 screen aimed at identifying genetic factors that contribute to breast cancer reactivation in DCCs. Here we demonstrate *mediator complex subunit 4* (*Med4*) as a novel cancer cell intrinsic gatekeeper in metastatic reactivation.

The Mediator complex regulates Pol II-mediated transcription by adjoining enhancer-bound transcription factors and the general transcription machinery in *cis*(El Khattabi et al., 2019; Whyte et al., 2013). Mediator, comprising a 26-subunit core organized into head, middle, and tail modules, and a four-subunit dissociable kinase module, manifests functional adaptability due to the variable arrangement of its subunits(Abdella et al., 2021; Verger et al., 2019). The subunit-specific composition and function of Mediator in its interaction with transcription factors (TF) imply that individual subunits of the complex regulate distinct transcriptional responses(Allen and Taatjes, 2015; Soutourina, 2018). The molecular basis for the disparate functional roles of individual subunits in various biological contexts remains incompletely understood(Barbieri et al., 2012; Ito et al., 2000; Stevens et al., 2002), particularly as it pertains to *Med4*. We show that genetic perturbation of *Med4* awakens dormant tumor cells in pre-metastatic niches. The clinical relevance of our findings is supported by the observation that low *MED4* expression is associated with increased risk of metastatic relapse in patients with breast cancer. Taken together, we propose a novel epigenetic regulatory mechanism whereby *Med4* depletion rewires integrin signaling through decompaction of 3-dimensional chromatin architecture and enhancer reprogramming.

## Results

### Genome-scale CRISPR KO screen identifies *Med4* as a dormancy regulator

To identify novel regulators enforcing breast cancer dormancy, we performed a genome-wide CRISPR knockout (KO) screen using the GeCKO v2 library containing 67,405 sgRNAs that was designed to target 20,611 distinct genes in the mouse genome (Gao et al., 2014; Joung et al., 2017) (Figure 1A). Following tail vein injection and lung infiltration, 4T07 breast cancer cells acquired the ability to exit dormancy, underwent reactivation and generated macrometastases (Figure S1A). After isolating tumor cells from each lung metastasis, we cloned and performed Sanger sequencing on the integrated sgRNA inserts to identify single genes that can reactivate metastasis and identified 26 unique metastases associated with a single sgRNA, which was discernible through Sanger sequencing (Table S1). Intriguingly, we identified sgRNAs targeting 3 genes (*Med4*, *Nipbl; Nipped-B-like protein*, and *Esco1; Establishment of sister chromatid cohesion N-acetyltransferase 1*) involved in the formation of super-enhancers among the top 3 hits from the initial validation. Whereas MED4 is a core subunit of the Mediator complex, which connects super-enhancers and promoters in *cis*, NIPBL and ESCO1 aid in loading cohesin onto DNA and hence physically stabilize the assembly of the super-enhancer-promoter complex (Figure S1B). We further examined the effects of knocking down each gene using multiple shRNAs in both 4T07-TGL and D2A1-d-TGL cells. *Med4* emerged as the most promising hit, showing biological potency across three different shRNAs and in both cell lines. (Figure 1B).

**Figure 1.**
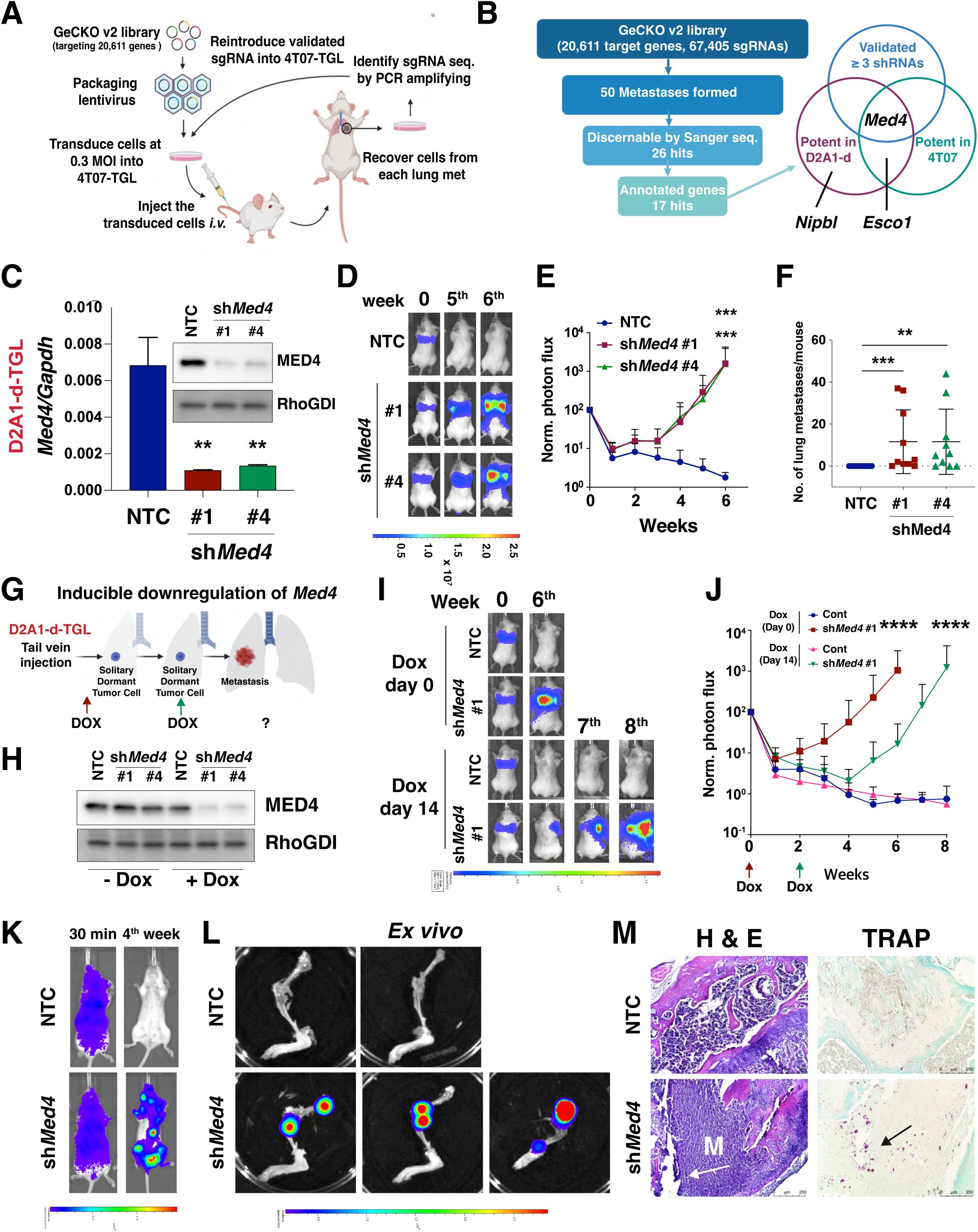
*Med4* was identified as a gatekeeper for metastatic reactivation by forward genetic screening using CRISPR-Cas9 system. **(A)** Schematic representation of the *in vivo* CRISPR-Cas9 screening strategy. **(B)** Schematic overview of the screening and validation process. The flow diagram outlines the progressive filtering and validation steps leading to the identification of the most promising target, Med4. **(C),** RT-qPCR analysis and immunoblots of *Med4* expression in D2A1-d-TGL cells expressing MirE shRNA targeting either non-targeting control (NTC) or *Med4.* RhoGDI was served as the loading control.; Mean values (± S.D.), p values; unpaired Student’s t-test. **(D-F),** D2A1-d-TGL cells transduced with either NTC or *Med4* shRNAs were inoculated *i.v.* in syngeneic mice. Lung metastasis was measured by bioluminescent (BLI) imaging at the indicated time. The panels show representative images (D), the graphs of the normalized BLI photon flux (E), and the graphs of the counts of macrometastases (F). Mean values (± S.D.), p values; Mann-Whitney U test (E and F). **(G-J)** Schematic representation of inducible expression experiment (G), Immunoblot analysis of MED4 in D2A1-d-TGL cells transduced with doxycycline (Dox)-inducible MirE shRNA targeting NTC or *Med4*. RhoGDI was served as the loading control. (H), D2A1-d-TGL cells expressing Dox-inducible MirE shRNA targeting NTC or *Med4* were inoculated *i.v.* in syngeneic mice. Dox was administrated either immediately after injection (Dox day 0) or 14 days after inoculation of the cells (Dox day 14). Lung metastasis was measured by BLI normalized photon flux at the indicated time. The panels show representative images (I), and the graphs of the normalized BLI photon flux (J); Mean values (± S.D.) p values; Mann-Whitney U test. **(K-M)** D2A1-d-TGL cells expressing shRNA targeting either non-targeting control (NTC) or *Med4* were inoculated intracardially (i.c.) in syngeneic mice. Representative BLI images of mice at indicated time points (K) and *ex vivo* BLI images of forelimb and hind limb bones (L) were captured. Bone sections were subjected to H&E staining and TRAP staining (M), with arrows indicating osteolytic bone lesions.

To confirm that the *Med4* gene was genuinely edited to promote metastasis, we performed indel PCR analysis and western blotting in the clone originally isolated from the screen. The results indicated that the clone which contains a sgRNA targeting *Med4*, carried heterozygous instead of homozygous deletion (one allele of the clone carries a nonsense mutation) (Figure S1C-S1F). We hypothesized that *Med4* is an essential gene with examination of the DepMap database supports dependency on *Med4* (Figure S1G)(Tsherniak et al., 2017). In agreement with the essentiality of *Med4*, we were not able to generate *Med4* KO clones presumably because of strong counterselection. Hereafter, we focused on *Med4* due to its strong selection in our genome-scale screen, its potent metastatic modulatory activity *in vivo*, its central role in gene transcriptional regulation, its genomic alteration in human breast cancer metastases (see below), and its unexplored role in governing metastatic dormancy.

### *Med4* enforces dormancy

To validate the biological potency of *Med4 in vivo*, we used MirE shRNA to target *Med4* in 4T07-TGL cells (Figure S1H-S1I) and another model for breast cancer dormancy, D2A1-d-TGL cells (Figure 1C-1F). Upon *Med4* silencing, the cancer cells acquired the capacity to efficiently colonize the lung when intravenously injected into immunocompetent mice. To determine if *Med4* silencing specifically induces reactivation from dormancy in early metastatic niches, we used a doxycycline-inducible promoter to express shRNAs targeting *Med4*. Mice were treated with doxycycline either immediately or 2 weeks after tail vein injection (Figure 1G). Whereas immediate knockdown of *Med4* induced metastatic outgrowth at around 2 weeks, a 2-week delay in *Med4* knockdown caused a similar delay in metastatic outgrowth (Figure 1H-1J), confirming that *Med4* downregulation re-awakens breast cancer cells from dormancy. Clinically, breast exhibits a propensity for metastasis to distinct organs, including the lungs, bones, brain, and liver(Obenauf and Massague, 2015). To assess the impact of *Med4* depletion on promoting colonization in other organs, we systemically administered cancer cells by injecting them into the left ventricle. While control cells remained dormant after 4 weeks, *Med4*-depleted cells exhibited a diffuse whole-body bioluminescence signal (Figure 1K). Subsequent dissection and *ex vivo* bioluminescence analysis revealed that metastases were exclusive to the bone following left ventricle inoculation, particularly in the epiphysis of the femur, tibia, and humerus (Figure 1L). Histological examination using H&E staining confirmed the presence of cancer cells within the epiphysis, along with associated bone lesions. Further analysis by TRAP staining identified the presence of osteoclasts near the lesions, indicative of osteolytic metastasis (Figure 1M). Taken together, these findings suggest that *Med4* maintains dormancy in breast cancer.

### *Med4* preserves the transcriptional and epigenetic landscape, preventing the formation of desmoplastic niches

To obtain a comprehensive view of the transcriptional reprograming in dormancy and metastatic reactivation, we performed bulk RNA sequencing (RNA-seq) of D2A1 (metastatic) cells and its dormant variant D2A1-d cells(De Cock et al., 2016). We identified a D2A1_metastatic gene signature (225 differentially upregulated genes in D2A1) and a D2A1_dormant gene signature (207 differentially upregulated genes in D2A1-d) (Figure S2A). We employed both signatures in analyzing bulk RNA-seq data from D2A1-d cells expressing two different shRNAs targeting *Med4*, in comparison to a control shRNA. Gene set enrichment analysis (GSEA) indicates that the D2A1_metastatic signature was enriched in *Med4*-silenced cells while the D2A1_dormant signature was enriched in control cells (Figure S2B). These results indicate that *Med4* depletion reprograms transcriptional profile from a dormant to a metastatic state. In addition, RNA-seq revealed that *Med4* silencing skews the transcriptomic profiles towards upregulation of gene expression, in contrast to the well-known function of the Mediator complex in transcription(Soutourina, 2018) (Figure 2A; Table S2-S5). To determine if the changes in the transcriptome resulting from *Med4* silencing are a general feature of the Mediator complex or specific to *Med4*, we employed MirE shRNA to target the Mediator complex subunit 1 (*Med1*) (Figure S2C). MED1 has been the most extensively investigated among the Mediator subunits and is recognized as a prominent transcriptional coactivator and a constituent of super-enhancers (SEs)(Hnisz et al., 2013; Whyte et al., 2013). Intriguingly, silencing *Med1* resulted in a skewed transcriptional profile towards repression which contrasts with the effect observed with *Med4* silencing (Figure S2D; Table S6-S8). GSEA using metastatic and dormant signatures also indicated that *Med1* reprograms the transcriptional profile in the opposite direction compared to *Med4* (Figure S2E).

**Figure 2.**
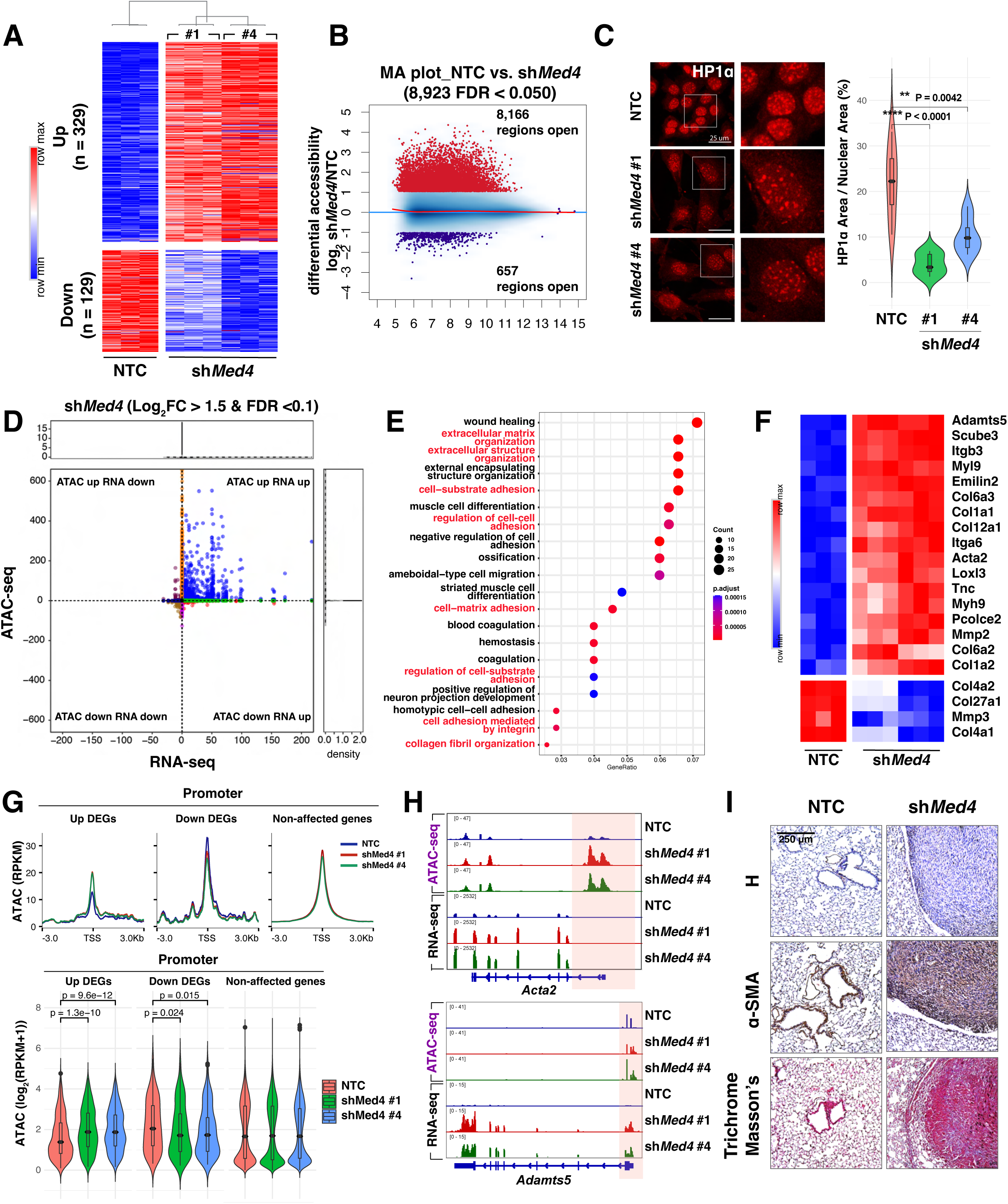
*Med4* inhibits the reprogramming of the transcriptional and epigenetic landscape to suppress matrix stiffening. **(A)** Heatmap of differentially expressed genes (DEGs; |Fold change| ≥ 2, FDR < 0.05) in NTC versus *Med4*-silenced D2A1-d-TGL one week after transduction. **(B)** Differential accessibility (log_2_ fold change in reads per accessible region) plotted against the mean reads per region. **(C)** D2A1-d cells expressing either NTC or *Med4* shRNAs were stained with HP1α (red). The panels display representative images (left) and a violin plot showing the ratio of heterochromatin foci puncta area to nuclear area (right). The data were obtained by measuring the area of heterochromatin puncta and nuclear area using ImageJ. Statistical significance was assessed using an unpaired t-test. **(D)** Intersection of RNA-seq and ATAC-seq plotted as - log_10_(FDR)*sign of the log_2_ fold change of *Med4*-sileced D2A1-d-TGL cells compared to NTC and each point representing one ATAC peak. **(E)** Gene ontology (GO) analysis of the genes located in the upper quadrant of (D) from the intersection of RNA-seq and ATAC-seq. **(F)** Heatmap shows the expression level of matrisome genes amongst the top differentially expressed genes in NTC versus *Med4*-depleted cells. **(G)** Metagene plots displaying ATAC-seq signal at promoter regions (around transcription start sites (TSS) within ±3 kb) for upregulated DEGs (Up DEGs), downregulated DEGs (Down DEGs), and non-affected genes (top). Violin plots showing the distribution of RPKM values for chromatin accessibility at promoter regions. Wilcoxon rank-sum test was used for statistical analysis (bottom). **(H)** IGV tracks showing ATAC-seq and RNA-seq profiles for on the *Acta2* and *Adamts5* genes in NTC and *Med4*-depleted D2A1-d-TGL cells. The promoter regions are highlighted in red. **(I)** Immunohistochemistry for α-SMA and Masson’s trichrome staining on lung metastases in mice that were intravenously injected with D2A1-d-TGL cells transduced with either non-targeting control (NTC) or *Med4* shRNAs.

We then investigated the impact of *Med4* depletion on the 3D chromatin structure. ATAC-seq analysis confirmed that *Med4* depletion led to a substantial increase in chromatin accessibility (Figure 2B). Conversely, *Med1* silencing had minimal impact on chromatin accessibility (Figure S2F). Complementary micrococcal nuclease (MNase) digestion assay(Mieczkowski et al., 2016) aligned with the findings from ATAC-seq results, showing that *Med4* depletion led to decompaction, particularly of poly-nucleosome part, indicative of heterochromatin structures (Figure S2G). On the contrary, *Med1* silencing resulted in increased resistance to MNase, suggesting that the downregulation of *Med1* caused chromatin structure to become more compact (Figure S2H). To directly observe the impact of *Med4* depletion on 3D chromatin architecture, we visualized chromatin using HP1α staining, a marker for constitutive heterochromatin(Wreggett et al., 1994). Intriguingly, *Med4*-depleted cells displayed a sparse distribution of heterochromatin foci accompanied by an increase in nuclear size, whereas the control cells exhibited a condensed heterochromatin structure (Figure 2C, and Supplementary Movies 1-3). In contrast, *Med1*-depleted cells did not exhibit significant changes in the distribution of heterochromatin foci (Figure S2I, and Supplementary Movies 4-6), highlighting *Med4*’s distinct role in the spatial organization of chromatin compared to *Med1*.

To delve deeper into the consequences of *Med4* silencing on chromatin structure and transcriptional output, we integrated RNA-seq and ATAC-seq results. Upon depleting *Med4*, the majority of genes exhibited significant changes in chromatin accessibility and gene expression, with increased levels in both accessibility and transcription (Figure 2D). Gene ontology (GO) analyses of genes showing concomitant changes in accessibility and gene expression (located in the upper quadrant of the intersection between RNA-seq and ATAC-seq) indicated significant enrichment of ECM-related and cell-adhesion-related gene signatures (Figure 2E). In particular, the majority of differentially expressed genes (DEGs) after *Med4* silencing were collagen-(e.g., *Col1a1, Col1a2, Col6a1*), actomyosin-related (e.g., *Acta2, Myl9*), and ECM-remodeling genes (e.g*., Loxl3, Mmp2, Adamts5*) (Figure 2F).

To confirm the link between chromatin structure and ECM gene expression regulated by *Med4*, further profiling of genome-wide chromatin accessibility revealed that *Med4* downregulation alters the promoter regions of DEGs, with upregulated DEGs exhibiting increased accessibility and downregulated DEGs showing reduced promoter accessibility (Figure 2G and Figure S2J). This suggests that *Med4* selectively modulates promoter chromatin accessibility, thereby influencing gene expression in a context-dependent manner, particularly in the regulation of ECM genes (Figure 2H).

Finally, this modulation of chromatin structure by *Med4* depletion is associated with stiffening matrices surrounding cancer cells, so called desmoplasia, which is a critical manifestation of cancer progression(Di Martino et al., 2022; Ishihara and Haga, 2022; Paszek et al., 2005; Ueno et al., 2015). Immunohistochemistry (IHC) of α-SMA (a protein coded by *Acta2*) and Masson’s trichrome stain, which labels collagen, validated the substantial desmoplastic stromal reaction in metastases induced by *Med4* silencing compared to the dormant niche (Figure 2I). Collectively, these results indicate that *Med4* haploinsufficiency bolsters matrix stiffening, which creates a favorable environment conducive to the metastatic outgrowth.

### *Med4* restrains enhancer priming and activation

We next explored how this chromatin structural change translates into broader epigenomic reprogramming. Histone modifications, often found in recurring combinations at promoters, enhancers, and repressed regions, serve as crucial markers for gene regulation (Barski et al., 2007; Ernst and Kellis, 2010, 2012). We performed ChIP-sequencing for promoters (H3K4me3), enhancers (H3K27Ac and H3K4me1), active transcription (H3K79me2), and heterochromatin (H3K9me3). Notably, *Med4* depletion led to a marked increase in these histone modifications particularly those associated with transcriptional activation (Figure S3A-S3E). To further dissect the epigenomic reprogramming, we employed the ChromHMM algorithm to model combinatorial patterns of the five histone modifications (Figure 3A). The resulting 10-state ChromHMM model categorized various epigenomic states, including promoters (E03 and E04), distal enhancers (E05, E06, and E07), genic or transcribed enhancers (E08 and E09), transcription (E10), and heterochromatin (E02). *Med4* silencing led to an increase in active histone modifications in promoter and enhancer regions, but slight decrease in heterochromatin regions (Figure S3F-S3I). GO analyses of these chromatin states revealed that distal enhancer states were associated with ECM gene signatures, consistent with our RNA-seq and ATAC-seq results (Figure 3B, and Figure S3J). We thereby focused on distal enhancer states, which were subdivided by three categories: active enhancers (E06), characterized by the enrichment of both H3K27Ac and H3K4me1; poised (closed) or primed (open) enhancers (E07), marked by H3K4me1 alone, reflecting regions primed for future activation; and weak enhancers (E05), marked by H3K27Ac alone. *Med4* depletion led to an increase in both H3K4me1 and H3K27Ac deposition in active and poised enhancers with a more robust increase in H3K4me1 (Figure 3C). Integration with ATAC-seq data revealed that *Med4* depletion opens up poised enhancer regions, transitioning them to a more accessible, primed state, ready for activation upon receiving the necessary cues (Figure 3D).

**Figure 3.**
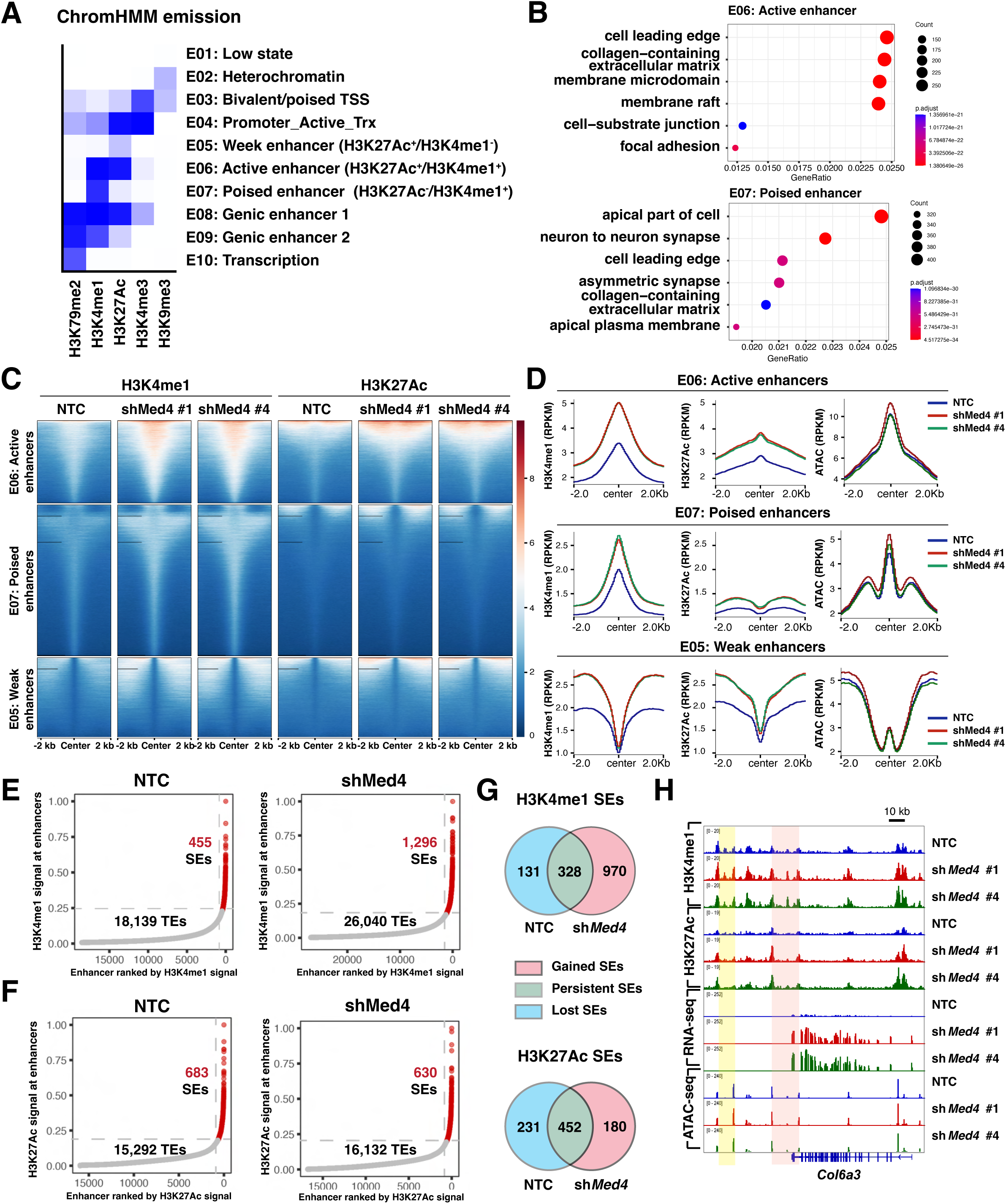
*Med4* maintains enhancer poise, restricting abnormal ECM gene expression. **(A)** Emission probabilities of the 10-state ChromHMM model on the basis of ChIP-seq profiles of five histone marks (shown in x axis) profiles in D2A1-d cells expressing MirE shRNA targeting *Med4* or NTC. Each row represents one chromatin state, and each column corresponds to one chromatin mark. The 10 states predicted using ChromHMM represent various enhancer states (E05, E06, E07, E08, and E09), promoter state (E03 and E04), transcription states (E10), and heterochromatin state (E02). Each column corresponds to a histone modification. **(B)** The top GO cellular component terms in active enhancer (E06; upper) states and poised enhancer (E07; bottom). **(C)** Heatmaps showing H3K4me1 and H3K27Ac across the individual peaks of distal enhancer chromatin states **(D)** Metagene plot of mean ChIP-seq signal of H3K4me1, H3K27ac and ATAC-seq signal across the individual peaks of each mark. Metagene analysis was centered on the middle of peaks and 4 kb around peak centers are displayed (2 kb upstream and 2 kb downstream). **(E)** Distribution of normalized H3K4me1 signal enrichment across enhancer regions in NTC (left) and Med4-silenced cells (right). A subset of regions with exponentially higher levels of H3K4me1 comprises super-enhancers (SEs; red), while the remainder are typical enhancers (TEs; grey). **(F)** The distribution of normalized H3K27Ac signal enrichment across enhancer regions in NTC (left) and Med4-silenced cells (right) similarly identifies SEs (red) and TEs (grey). **(G)** Intersection of H3K4me1-enriched SEs (left) and H3K27Ac-enriched SEs (right) between NTC and shMed4 cells. **(H)** Genome tracks showing H3K4me1-enriched SEs associated with the *Col6a3* gene. The *de novo* SE in sh*Med4* is highlighted in red and the persistent SE is highlighted in yellow.

Enhancers are classified into super-enhancers (SEs) and typical enhancers (TEs). SEs, which are clusters of enhancers with high levels of transcriptional co-activators, are crucial for regulating key genes involved in cell identity and disease(Hnisz et al., 2013; Whyte et al., 2013). To explore the epigenomic reprogramming induced by *Med4* depletion, we extended our analysis to include the identification and characterization of SEs. Our analysis showed that *Med4* depletion increases the intensities of both H3K4me1 and H3K27Ac at their respective SE and TE regions (Figure S4A-S4D). Interestingly, *Med4* silencing led to a significant increase in the number of H3K4me1-enriched SEs (455 in NTC versus 1,296 in shMed4) and TEs (18,139 in NTC versus 26,040 in shMed4) compared to the H3K27Ac-enriched SEs (683 in NTC versus 630 in shMed4) and TEs (15,292 in NTC versus 16,132 in shMed4) (Figure 3E and 3F). The intersection of SEs between NTC and *Med4*-silenced cells reveals a substantial increase in *de novo* H3K4me1-enriched SEs (Figure 3G), which are associated with cell-cell adhesion, focal adhesion, and collagen-related ECM (Figure S4E and S4F). While the number of *de novo* H3K27Ac-enriched SEs did not significantly change compared to H3K4me1-enriched SEs, the intensity of H3K27Ac marks increased significantly in *Med4*-depleted cells, suggesting that *Med4* depletion contributes to the full activation of SEs. The *Col6a3* gene exhibits both a *de novo* H3K4me1-enriched SE and a persistent SE, each with significantly higher H3K4me1and H3K27Ac intensities compared to the control (Figure 3H). Collectively, these observations suggest that *Med4* inhibits H3K4me1 deposition, preventing the transition of enhancers from a poised to an active state. This inhibition restricts epigenomic reprogramming and ECM gene expression.

### *Med4* suppresses cancer stemness and expansion in a desmoplastic niche

Given the functional similarities between metastasis-initiating cells and cancer stem cells (CSCs) (Malanchi et al., 2011), we hypothesized that *Med4* might regulate CSC traits. Indeed, GSEA showed enrichment of the Wnt pathway in *Med4*-depleted cells, a pathway known to be associated with cancer stemness (Figure S5A) (Reya and Clevers, 2005). *Med4*-depleted cells exhibited a substantial increase in their self-renewal capacities, demonstrated by the increased number and size of tumor spheres (Figure S5B and S6A), enhanced clonogenicity (Figure S5C and S6B), and improved anchorage-independent growth (Figure S5D and S6C). In addition, *Med4*-depleted cells exhibited increased migratory (Figure S5E and S6D) and invasive (Figure S5F) capacities which are common characteristics of cells undergoing epithelial to mesenchymal transition (EMT)(Pastushenko and Blanpain, 2019). Overall, these results suggest that *Med4* depletion endows dormant cancer cells with self-renewal capacities.

As CSCs exhibit preferential growth in a stiff matrix, a hallmark of the tumor microenvironment, this suggests that mechanical cues from the ECM can synergize with intrinsic stemness to promote cancer cell metastatic abilities (Li et al., 2023; Liu et al., 2023). Thus, we sought to examine if *Med4* mediates outgrowth in response to the stiffness and composition of matrices. To model a normal ductal microenvironment characterized by a basement membrane-rich ECM, and a stiff tumor microenvironment enriched in collagen I(Egeblad et al., 2010; Paszek et al., 2005), we placed control and *Med4*-silenced breast cancer cells in two different conditions: (1) 3D Matrigel, representing the basement membrane-rich ECM, and (2) 3D Matrigel plus collagen I, which recapitulates the tumor microenvironment. Strikingly, *Med4* depletion promoted tumor organoid growth and invasion in stiff 3D matrices (Matrigel + collagen I), while cells maintained a spherical growth pattern in the soft 3D matrices (Matrigel-only) regardless of their *Med4* expression (Figure 4A). These data indicate that cells with a *Med4*-haploinsufficient background can preferentially grow in the desmoplastic niche. Collectively, these results indicate that *Med4* haploinsufficiency not only induces matrix stiffening, but also stimulates the cancer-cell intrinsic survival and proliferative abilities within the established metastatic niche.

**Figure 4.**
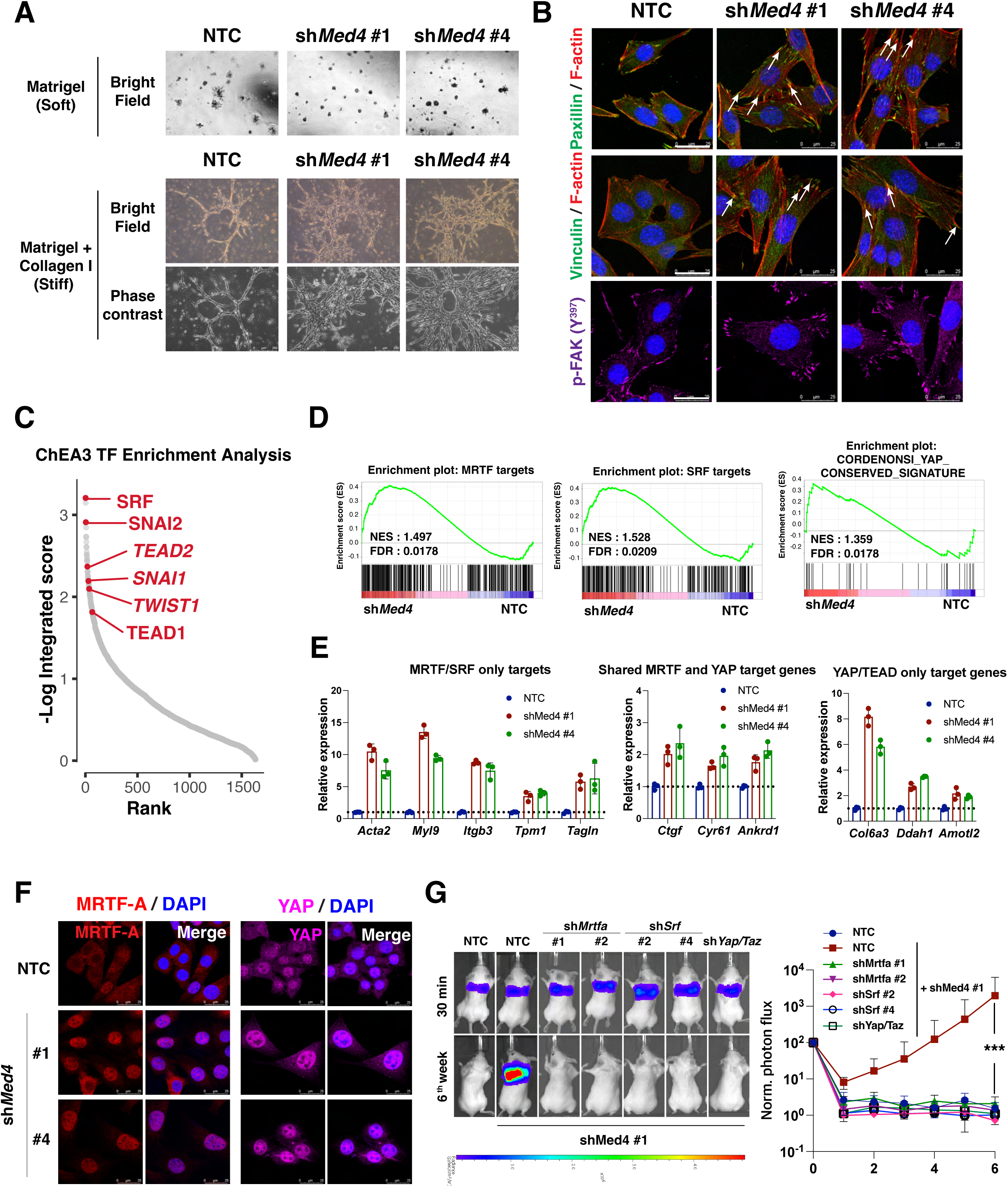
*Med4* sustains dormancy by suppressing integrin-mediated mechano-transduction. **(A)** D2A1-d cells expressing MirE shRNA targeting *Med4* or NTC were cultured on 3D Matrigel with or without collagen I. **(B)** D2A1-d cells expressing MirE shRNA targeting Med4 cultured on 2D were stained for either vinculin or paxillin (green), F-actin (red), p-FAK (purple) and the nuclei were counterstained with DAPI (blue). Arrows indicate the focal adhesion plaques. **(C)** ChEA3 TF enrichment analysis of the promoters of the genes upregulated in *Med4*-depleted cells compared to NTC. **(D)** Gene signature enrichment analysis (GSEA) of MRTF, SRF, and YAP transcriptional signatures. NES, normalized enrichment score; FDR, false discovery rate (n = 3 biological replicates per condition). **(E)** RT-qPCR results were indicated as relative mean expression of MRTF/SRF only target genes, YAP only target genes and shared target genes in either control or *Med4*-depleted D2A1-d cells. **(F)** D2A1-d cells expressing MirE shRNA targeting *Med4* cultured on 2D were stained for MRTF-A (red), or for YAP (purple). The nuclei were counterstained with DAPI (blue). **(G)** D2A1-d-TGL cells co-transduced with indicated MirE shRNAs were inoculated *i.v.* in syngeneic mice. Lung metastasis was measured by bioluminescent (BLI) imaging at the indicated time. The panels show representative images (left), and the graphs of the normalized BLI photon flux (right). Mean values (± S.D.) p values; Mann-Whitney U test.

### *Med4* suppresses integrin-mediated nuclear mechanosensing

To exit dormancy, metastatic cancer cells extend lamellipodia and establish productive interactions with the matrix. Adhesion molecules, particularly integrins, play a crucial role in transmitting external stimuli from the ECM(Barkan et al., 2010; Case and Waterman, 2015; Kechagia et al., 2019). Considering the transcriptional alterations in the ECM composition (Figure 2E) and integrin repertoire (Figure S7A) by *Med4* depletion, we hypothesized that *Med4* haploinsufficiency promotes metastatic reactivation by activating integrin-mediated signaling. Immunofluorescence analysis revealed that *Med4*-silenced cells exhibit a more organized actin cytoskeleton and more numerous focal adhesion plaques (Figure 4B and S7B). Our data also show that phosphorylation of key integrin signaling molecules was significantly increased upon *Med4* depletion (Figure S7C). Collectively, these findings provide evidence that *Med4* inhibits integrin signaling activation.

To identify the top transcription factors controlling the transcriptome changes by *Med4* silencing, we applied ChIP-X Enrichment Analysis 3 (ChEA3) transcription factor enrichment analysis to the RNA-seq data(Keenan et al., 2019). Notably, we found that SRF, and TEAD 1/2 cistromes govern the transcriptional reprogram by *Med4* depletion along with EMT cistromes (Figure 4C). Also, *de novo* motif enrichment analysis performed on genes exhibiting concurrent increases in accessibility and gene expression following *Med4* silencing confirmed the enrichment of TEAD, TEAD4, and SRF motifs (Figure S8A). SRF, a co-factor for myocardin-related transcription factor (MRTF), and TEADs, transcriptional co-activators for YAP and TAZ, are major mechanosensitive transcriptional factors that transduce mechanical cues by the cytoskeleton to gene expression in response to integrin-mediated adhesion. GSEA results revealed that the gene signatures for MRTF/SRF(Esnault et al., 2014) and for YAP(Cordenonsi et al., 2011) are enriched in *Med4*-depleted cells (Figure 4D). In response to both biochemical and mechanical signals, YAP undergoes translocation from the cytoplasm to the nucleus, where it interacts with and triggers the activation of the TEAD transcription factor(Zhao et al., 2008). MRTF binds stoichiometrically to soluble G-actin and is released as a consequence of actin polymerization and formation of stress fibers. Once released from G-actin, MRTF enters into the nucleus, where it functions as a co-factor for SRF, which drives expression of cytoskeletal genes in a feed forward loop(Posern and Treisman, 2006). Hence, we sought to investigate whether mechanosensitive transcription factors, specifically MRTF and YAP, were activated upon *Med4* depletion. We observed activation of MRTF/SRF target genes(Calvo et al., 2013; Foster et al., 2017), YAP/TEAD target genes(Dupont et al., 2011), and shared MRTF and YAP target genes in *Med4*-depleted cells (Figure 4E). Immunofluorescence results revealed that *Med4* silencing leads to nuclear localization of both MRTF-A and YAP (Figure 4F). To address the discrepancy between MRTF and YAP activation, we conducted time-course experiments using a doxycycline-inducible promoter for *Med4* shRNAs (Figure S8B-S8F). Our findings demonstrated that mRNA levels of MRTF/SRF target genes were significantly upregulated at 24 hours post-, whereas activation of the YAP/TEAD pathway was less robust and occurred 3 to 5 days later, prompting us to conclude that *Med4* depletion activates MRTF to a greater degree than YAP. Restoring MED4 expression by overexpressing *MED4* cDNA led to the relocation of MRTF-A to the cytoplasm and resulted in the reversal of MRTF-A/SRF target gene expression (Figure S8G and S8H). To determine if these integrin-mediated mechanosensitive transcription factors play a pivotal role for *in vivo* metastatic reactivation by *Med4* depletion, we depleted these mechanosensitive transcription factors though shRNA silencing and injected cancer cells into immunocompetent BALB/c recipients (Figure S8I). Silencing *Mrtfa*, *Srf,* or *Yap* + *Taz* abrogated the metastatic capacity of *Med4*-silenced cells, pointing to a requirement for these transcriptional regulators in driving metastatic reactivation (Figure 4G). Together, these findings indicate that *Med4* restrains the initiation of integrin-mediated mechanotransduction to maintain dormant states.

### *MED4* haploinsufficiency correlates with poorer prognosis in breast cancer

To explore the clinical relevance of *MED4* status in breast cancer patients, we examined *MED4* expression and prognosis in multiple metastatic breast cancer datasets from cBioportal(Cerami et al., 2012; Gao et al., 2013). *MED4* exhibited shallow deletions in almost 40% of human breast cancer metastasis samples (Figure 5A, Figure S9A and S9B)(Lefebvre et al., 2016). The low proportion of deep deletions further underscores *MED4* as an essential gene. Consistent with genomic analysis, paired RNA-seq revealed that mRNA levels of *MED4* are lower but not undetectable in lung, liver, and brain metastases than in matched primary tumors (Figure 5B)(Siegel et al., 2018). Intriguingly, Kaplan-Meier analysis of primary tumor datasets revealed that *MED4* expression levels could be used to stratify patients into high- and low-risk groups in terms of recurrence-free survival; lower *MED4* levels in primary tumors were strongly correlated with poor prognosis in patients with advanced breast cancer across subtypes (Figure 5C and Figure S9C)(Gyorffy et al., 2010). Furthermore, lower levels of *MED4* mRNA can serve as a prognostic indicator for poor overall survival, particularly in subgroups of patients with estrogen receptor-negative (ER-) or human epidermal growth factor 2-positive (HER2+) breast cancer (Figure S9D and S9E). We confirmed the prognostic value of *MED4* gene via SurvExpress platform. *MED4* could effectively stratify patients into high and low-risk groups for distant metastasis-free survival (DMFS), with lower *MED4* expression in high-risk group (Figure 5D). Finally, the *MED4*-associated ECM gene signature (Figure 2F) was found to be associated with DMFS, further validating its predictive capability (Figure 5E). Collectively, these results suggest that *MED4* and its associated gene signatures could be prognostic markers to predict metastatic relapse in breast cancer.

**Figure 5.**
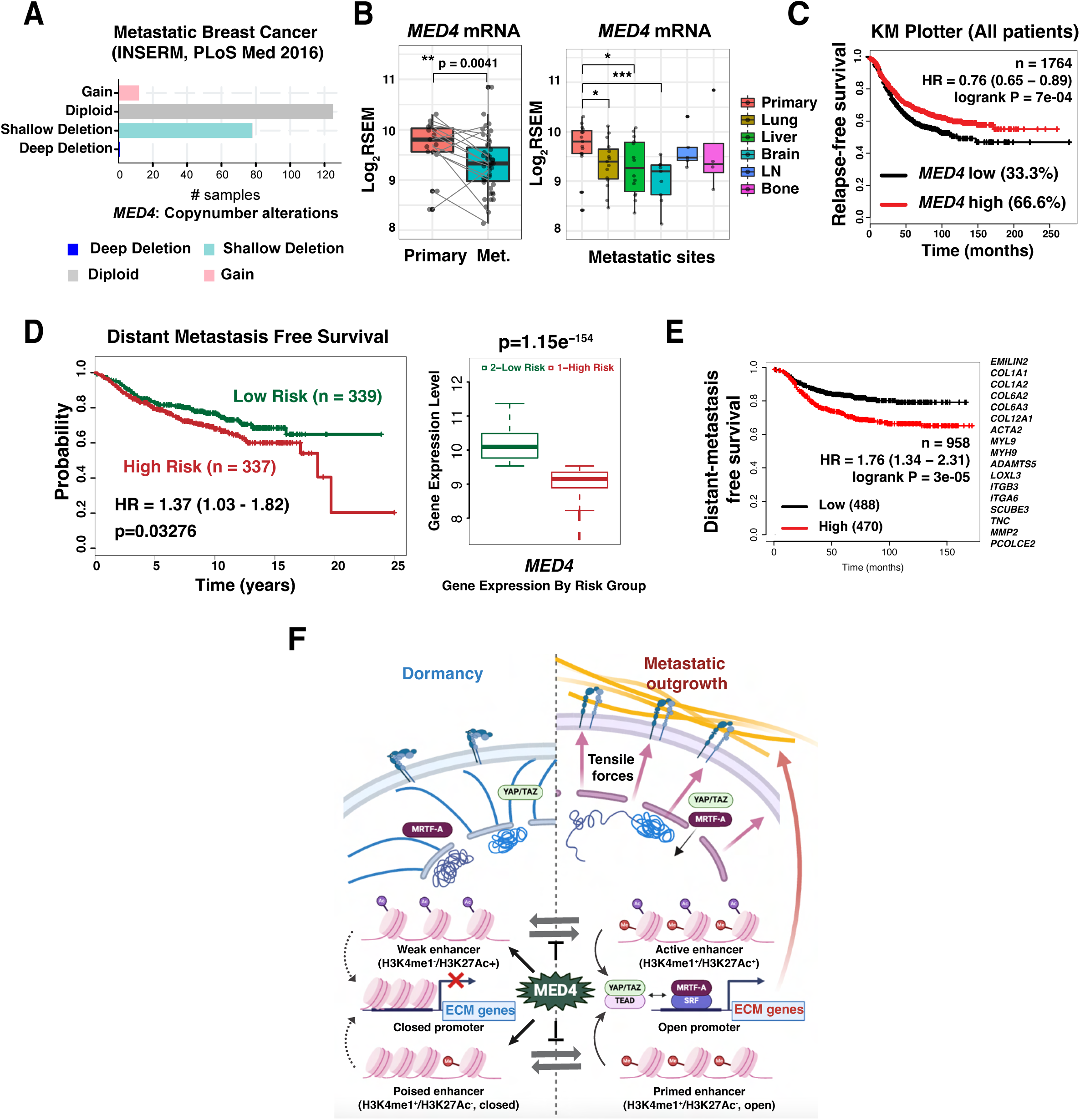
*MED4* haploinsufficiency and associated haplo-signatures are correlated with metastatic relapse in patients. **(A)** Copy number alterations in metastatic breast cancer patients (from INSERM) were analyzed by cBioportal database. **(B)** *MED4* mRNA levels were measured by paired RNA-seq of primary tumors and multiple metastases. **(C)** Kaplan-Meier analysis of relapse-free survival (RFS) of all patients in publicly available breast cancer datasets (source KM Plotter for breast cancer). **(D)** Patients were stratified according to *MED4* expression using a probe set, 222438_at as indicated. HR, hazard ratio. Kaplan-Meier curve of the Breast Cancer Meta-base: 10 cohorts 22K gene data showing the distant metastasis-free survival (DMFS) of patients separated in 2 cohorts based on the risk (left). Red and green curves denote high- and low-risk groups respectively. Box plot across risk groups, including the p-value testing for difference using t-test (right). **(E)** Kaplan-Meier analysis of DMFS in patients stratified by the expression of the *MED4*-associated haplo-gene signature. **(F)** Schematic illustration of MED4’s role in maintaining the epigenomic landscape and cellular dormancy. *Med4* inhibits enhancer activation (weak to active) and priming (poised to primed) by suppressing H3K4me1 deposition. *Med4* depletion disrupts this control, leading to epigenomic and transcriptional reprogramming that rewires ECM protein and integrin expression, promoting integrin-mediated mechanotransduction. Stress fiber assembly further reinforces chromatin modifications by exerting tension on the nuclear membrane. *Med4* acts as a gatekeeper of dormancy by regulating enhancer dynamics.

## Discussion

In this study, a genome-scale screen and functional *in vivo* assays identified and validated *Med4* as a critical gatekeeper of breast cancer metastasis dormancy. Mechanistically, we establish that *Med4* depletion causes decompaction of 3D chromatin structure and initiates a novel enhancer reprogramming event, wherein H3K4me1 deposition transforms weak and poised enhancers into more active states. This enhancer reprogramming unlocks ECM gene expression, activates integrin-mediated mechano-transduction, and facilitates the emergence of cancer cells from dormancy in metastatic niches. Correspondingly, clinico-genomic analyses reveal that heterozygosity and associated lower expression of *MED4* correlates with worse prognosis and decreased metastasis-free survival. Thus, this work highlights a previously unrecognized role of *Med4* haploinsufficiency in regulating enhancers and gene expression programs that promote functional tropism and metastatic reactivation (Figure 5F).

We demonstrate that *Med4* haploinsufficiency has the potential to initiate metastatic reactivation within the context of breast cancer. Interestingly, our analysis reveals that *Med4* silencing predominantly stimulates a transcriptional profile consisting of ECM-related genes, which stands in contrast to the Mediator’s conventional role in transcription. Conversely, the silencing of *Med1* displayed a unique and directionally oppositional function that MED4 serves within the context of Mediator-mediated transcription, highlighting the considerable flexibility of the complex contingent on its subunit composition. Sun and colleagues conducted a shRNA screening targeting individual subunits of transcription factor IID (TFIID) and the Mediator complex(Sun et al., 2021). Disrupting various subunits yielded distinct effects on the transcriptome of mouse embryonic stem cells (mESCs), even within subunits of the same complex. Their shRNA-based screen, similar to our findings, revealed that *Med4* shRNA altered transcriptome towards activation, and gene ontology analysis of the up-regulated gene list highlighted the enrichment of ECM signatures. However, when they employed the auxin degron system for the acute degradation of MED4, it had a minimal effect on the integrity of the Mediator complex and resulted in transcriptome downregulation by decreasing RNA polymerase II recruitment to the promoters. Hence, it is plausible that a dosage effect whereby *Med4* knockdown as opposed to complete knockout drives the pathological phenotype.

The Mediator complex regulates transcription by linking enhancers to promoters, with super-enhancers characterized by high Mediator binding and associated with key genes and oncogenes(Whyte et al., 2013). While the term ‘Mediator’ is often used as a general marker for defining SEs, it is commonly represented by only MED1 and MED12(Guo et al., 2019; Hnisz et al., 2013). To address the controversial role of *Med4* in transcriptional repression, we focused on its impact on epigenomic reprogramming. Our integrated analysis of RNA-seq, ATAC-seq, and histone ChIP-seq data reveals that *Med4* depletion leads to significant epigenomic reprogramming, making the genome more permissive to transcriptional activation. We found that distal enhancers marked by H3K4me1 and H3K27Ac play a crucial role in differential gene expression, consistent with the established role of super-enhancers in defining cell identity. Specifically, *Med4* knockdown leads to a notable increase in H3K4me1 deposition at the enhancer regions compared to H3K27Ac, and differential expression and chromatin accessibility were correlated with distal enhancer reprogramming. This suggests that *Med4* plays a significant role in keeping enhancers in a poised state, and its depletion shifts the balance toward preparing more enhancers for potential activation, thus contributing to epigenomic reprogramming and altered gene expression, particularly in ECM genes. Our results revealed that *Med4* selectively regulates chromatin accessibility at promoter regions of differentially expressed genes. Recent studies indicate that H3K4me1 facilitates enhancer-promoter interactions(Kubo et al., 2024), suggesting that *Med4* depletion may facilitate the interaction between activated and primed enhancers with opened promoter region. Further investigation using Hi-C or promoter capture Hi-C is needed to explore these enhancer-promoter interactions in detail. Additionally, biochemical studies are required to elucidate how *Med4* directly regulates H3K4 monomethylation, potentially involving H3K4 mono-methyltransferases such as KMT2C or KMT2D(Alam et al., 2020). Moreover, it is worth noting that *Med4* depletion led to a significant increase in nuclear size, which may contribute to the observed spatial reconfiguration in 3D chromatin structure. Given that the nuclear lamina is known to anchor heterochromatin to the nuclear periphery for gene silencing (Reddy et al., 2008; van Steensel and Belmont, 2017), future investigations into the impact of nuclear architecture changes could provide deeper insights into the mechanisms governing chromatin organization.

Current adjuvant treatment for breast cancer is aimed to prevention of metastatic relapse, and despite the prognostic value of pathological, genomic, and transcriptional testing, late metastasis occurs in nearly 25% of patients(Berry et al., 2005). Elucidating the molecular mechanisms that enable dormant tumor cells to evade the immune system and survive at early metastatic sites, as well as identifying subsets of patients who could benefit from therapy targeting these mechanisms may reduce the mortality related to metastatic relapse. We found that a large fraction of metastatic breast cancer samples from patients have shallow *MED4* deletions (Figure 5A, S9A and S9B), and lower *MED4* levels correlated with poor prognosis in patients with advanced breast cancer across subtypes (Figure 5C-5D and S9C-S9E), suggesting that *MED4* may play a critical role in governing metastatic dormancy in breast cancer patients.

Our findings highlight the potential of *MED4* expression and related gene signatures as a biomarker of metastatic dormancy. Future studies will investigate whether *MED4* can stratify patients prone to recurrence, enabling the identification of subsets at high risk for subclinical metastasis and may uniquely benefit from therapeutic intensification. Moreover, these data lay the foundational groundwork for novel strategies targeting effectors activated by *MED4* haploinsufficiency including inhibitors of the FAK or YAP/TEAD pathways.

## Materials and Methods

### Animal studies

Mouse studies were conducted in accordance with protocols approved by the Institutional Animal Care and Use Committee of Columbia University and MD Anderson Cancer Center. Six-week-old female mice BALB/c were purchased from The Jackson Laboratory.

To examine lung colonization, 4T07 (3.0 x 10^5^), D2A1-d (2.0 x 10^5^), and D2A1 (2.0 x 10^5^) mammary carcinoma cells were resuspended in 100 µl of PBS and injected into the lateral tail vein. Doxycycline (Dox) was administered to mice through water *ad libitum* (2 mg/ml). For inducible shRNA experiments, Dox was administered either immediately after injection (Dox day 0) of the cancer cells or 14 days later (Dox day 14). Bioluminescence (BLI) was used to examine metastatic outgrowth. Metastatic lesions were confirmed by postmortem dissection. To examine multi-organ colonization, D2A1-d-TGL (8.0 × 10^4^) cells were injected into the left ventricle of BALB/c mice in 100 µl of PBS.

For bioluminescent imaging, mice were anesthetized and injected retro-orbitally with 1.5 mg (75 mg/kg) of D-luciferin (Gold Biotechnology; LUCK-1G) at the indicated times after injecting cells. Animals were imaged in an IVIS Spectrum chamber within 5 min after D-luciferin injection, and data were recorded using Living Image software (PerkinElmer). To measure lung colonization, photon flux was calculated for each mouse by using a circular region of interest encompassing the thorax of the mouse. After subtracting a background value obtained from a control mouse injected only with D-luciferin, photon flux was normalized to the value obtained 30 min after injection of the tumor cells. This latter value was arbitrarily set at 100. For *ex vivo* analysis, anesthetized mice were injected with D-luciferin. Mice were euthanized 15 min later, and organs were rapidly dissected for analysis.

### Cell lines

The 4T07 cell line was generously provided by Dr. Fred Miller (Wayne State University, Detroit, MI) and cultured as described (Aslakson and Miller, 1992). The D2A1-d and D2A1 cell lines were generously provided by Dr. Robert Weinberg (MIT, Boston, MA) and cultured as described(De Cock et al., 2016). HEK293FT cells (from Thermo Fisher Scientific) was cultured per the manufacturer’s protocol. For bioluminescent tracking, cell lines were infected with a lentiviral vector encoding herpes simplex 3 virus thymidine kinase 1, green fluorescent protein (GFP) and firefly luciferase. The top 10% of GFP-positive cells were isolated by FACS to ensure uniform expression levels of GFP across all cell populations, mitigating any potential artificial effects caused by GFP’s immunogenicity.

### Generation of stable cell lines by lentiviral transduction

Stable cells were generated by lentiviral transduction as described in with a minor modification. In the case of constitutively expressing stable cells, all experiments were conducted within a 1 to 3-week window from transduction to minimize the counterselection effect of *Med4* depletion unless otherwise specified. In the case of cells expressing doxycycline-inducible constructs, the indicated time frame starts from the initiation of doxycycline treatment.

### Genome-wide CRISPR-Cas9 screening

A forward genetic screening was performed as described in with a minor modification(Gao et al., 2014). The GeCKO v2 library A was obtained from Addgene (1000000052) and was amplified and packaged in lentiviruses as described previously(Joung et al., 2017). The viral titer was determined by limiting dilution-colony counting. 4T07 cells were transduced with the library at the MOI of 0.3 to ensure one sgRNA/cell. Library-transduced cells were selected with 2 µg/ml of puromycin for 7 days, and then injected through tail vein of 6-week-old female BALB/c mice. Mice were monitored weekly by bioluminescent imaging and lung metastases were confirmed by postmortem dissection at 3-4 weeks after injecting library-transduced cells. Macroscopic lung nodules were dissected separately and digested in Collagenase Type IV (Stem Cell technologies; 07909) at 37°C for 1.5 h with rocking. After centrifuging at 1,000 rpm for 3 min, pellets were resuspended in the complete medium and filtered through 35 µm strainers. Isolated clonogenic tumor cells were expanded in 4 µg/ml of puromycin medium. Genomic DNAs were isolated using Maxwell^®^ RSC Cultured Cells DNA Kit (Promega; AS1620) according to the manufacturer’s protocol. Inserted sgRNA sequences were amplified by PCR with the following primers and thermocycling parameters (v2Adaptor-F: 5’-AATGGACTATCATATGCTTACCGTAACTTGAAAGTATTTCG-3’, v2Adaptor-R: 5’-TCTACTATTCTTTCCCCTGCACTGTTGTGGGCGATGTGCGCTCTG-3’; 95°C for 2 min, 30 cycles of (98°C for 1 s, 60°C for 5 s, 72°C for 35 s), and 72°C for 1 min)

All PCR was performed using KAPA HiFi HotStart ReadyMix (Kapa Biosystems; KK2602). The PCR products were isolated by QIAquick PCR Purification Kit (Qiagen; 28104) and subjected to Sanger sequencing to discern single entities that can reactivate metastasis. Consequently, we identified 26 metastases, each associated with a single sgRNA, which were discernible through Sanger sequencing. The table listing the gene names and corresponding sgRNA sequences is provided in **Table S1**. To validate hits, the sgRNAs were cloned in lentiCRISPR v2 vector, and transduced into 4T07-TGL cells. Transduced clones were injected through tail vein of 6-week-old female BALB/c mice and lung metastases were monitored weekly by bioluminescent imaging to prioritize hits.

### Indel analysis for validating a knockout screening

A pair of primers was designed to amplify the 250-300 bp region centered around the sgRNA cut site (GRCm39 chr.14 73752726-73752745). 300 ng of genomic DNA isolated in *Genome-wide CRISPR-Cas9 screening* was amplified by PCR with the following primers and thermocycling parameters (*Med4* indel primer_F: 5’-CACGGCCATGGCTTTCTAAC-3’, *Med4* indel primer_R: 5’-GAGGGTTGTGAGCTACGTGT-3’; 95°C for 2 min, 30 cycles of (98°C for 1 s, 60°C for 5 s, 72°C for 35 s), and 72°C for 1 min). PCR was performed using NEBNext® High-Fidelity 2X PCR Master Mix (NEB; M0541S). The purified PCR products were separated by a 4-20% TBE gel (Invitrogen; EC62252BOX) and stained with GelRed for 1 hr. The bands were visualized by ChemiDoc (Bio-Rad).

### DNA constructs for overexpression and silencing

The cDNA encoding human *MED4* was obtained from Addgene (15425). The cDNA was sub-cloned in FLAG-HA-pcDNA3.1-(Addgene; 52535) to generate N-terminally Flag-HA tagged MED4 cDNA. The N-term Flag-HA-MED4 cDNA was sub-cloned in pCDH-EF1-MCS-IRES-Neo (System Biosciences; CD502A-1) and verified by sequencing. The shRNAs were designed using the SplashRNA algorithm(Pelossof et al., 2017). shRNA sequences are shown in **Table S9**.

### Cell growth assay

Cells were seeded at a density of 1.0 x 10^5^ cells onto 6 cm dishes (D2A1-d cells) or 10 cm dishes (4T07 cells) at Day 0. Cells were trypsinized and counted at the indicated.

### Tumorsphere assay

Single cells suspension of 4T07, D2A1-d, and D2A1 cells (10,000 cells/ 2 ml) were plated on 6-well ultra-low attachment plates for tumorsphere cultures. Cells were cultured in serum free MEBM supplemented with 1:50 B27, 10 ng/ml bFGF (BD Biosciences;) and 20 ng/ml EGF (BD Biosciences;) and 4 ug/ml insulin, 4 ug/ml heparin, and 0.4% BSA for 10 days. Tumorsphere were visualized under phase contrast microscope, photographed and counted by GelCounter and represented graphically.

### Soft agar colony formation assay

6-well culture plates were coated with 1 ml of 0.6 % Low Melt Agarose (Bio-Rad; 1613111). Cells were diluted at a density of 1.0 x 10^4^ cells/ml in media containing 0.3% Low Melt Agarose and seeded 1 ml/ well. After 10 days, cells were stained with 0.005% crystal violet in 10% neutral buffered formalin (NBF) for 30 min at R.T. Soft agar was destained with distilled water until the background got clear. Colonies were scanned for visualization and counted manually.

### 3D culture

8-well chamber slides were coated with either 100 µl of 100% Matrigel (Corning; 354263) or 50% Matrigel + 50% Collagen I (Corning; 354236) at 37°C for 30 min prior to cell seeding (Nguyen-Ngoc et al., 2015). Cells were seeded onto the matrix at a density of 400 cells/100 µl/well.

### Clonogenicity assay

For measuring clonogenicity capacities, D2A1-d and 4T07 cells were seeded onto 10 cm dishes at a density of 1,000 cells. After 6 days, the morphology and size of colonies were captured by a phase-contrast microscopy. Cells were fixed with 100% methanol for 20 min and stained with 0.5% crystal violet in 20% methanol for 5 min. Then plates were destained with PBS until the background got clear. The images of whole plates were taken by a scanner and colonies were counted manually.

### Transwell migration/invasion assay

Migration assays were performed as previously described in (Bae et al., 2015) with minor modification. using uncoated cell culture inserts with 8 μm pores (Corning; 3422) according to manufacturer’s instructions. Briefly, D2A1-d and 4T07 cells were harvested with Trypsin/EDTA and resuspended in serum-free medium at a density of 1.0 x 10^5^ cells/ml for D2A1-d and 2.0 x 10^5^ cells/ml for 4T07. 100 µl of cells were seeded into the upper chamber of uncoated cell culture inserts with 8 μm pores (Corning; 3422). Lower chambers were filled with 650 µl medium containing 10% FBS as a chemoattractant. After 18 hours, cells were fixed and permeabilized with 70% ethanol for 15 min and stained with 0.1% crystal violet/10% ethanol for 20 min. The non-migrating cells on the upper surface of the filter were removed by cotton swab. The number of migratory cells was measured by counting at × 200 magnifications using a microscope. For invasion assays, Matrigel was diluted in ice-cold serum-free medium to a concentration of 300 µg/ml. 100 µl of diluted Matrigel solution was added into the upper chamber and incubated at 37°C for 30 min to coat the membrane. D2A1-d (1.0 x 10^4^ cells) and 4T07 (3.0 x 10^4^ cells) cells were seeded into the upper chamber and incubated for 24 hours. The following procedure was performed as described above.

### Micrococcal nuclease digestion assay (MNase digestion assay)

Global chromatin accessibility was assessed by MNase digestion assay as described previously(Cheng et al., 2019) with minor modification. 1 million cells were cross-linked with 1% formaldehyde for 10 min at R. T. and quenched with 125 mM glycine for 5 min. Cells were washed once with cold PBS and collected by centrifuging at 1,000 rpm for 5 min at 4°C. Fixed cells were re-suspended in lysis buffer (10 mM Tris-HCl, pH7.5, 10 mM NaCl, 0.5% NP-40) and incubated on ice for 10 min. After centrifugation, the cell pellets ware washed once with MNase digestion buffer (20 mM Tris-HCl, pH7.5, 15 mM NaCl, 60 mM KCl, 1 mM CaCl_2_). After re-suspended in 250 µl MNase digestion buffer with proteinase inhibitors, 62.5 units per 1 million cells MNase (NEB; M0247S) was added to resuspended pellets. After incubation at 37°C for the indicated time, 250 µl of STOP buffer (100 mM Tris-HCl, pH 8.1, 20 mM EDTA, 200 mM NaCl, 2% Triton X-100, 0.2% sodium deoxycholate) was added. Lysates were sonicated for 4 cycles (30 sec on and 30 sec off) in Bioruptor and supernatant was collected after centrifugation. The supernatant was reverse-crosslinked at 65°C overnight and treated with RNase A at 37°C for 1 h and Proteinase K at 55°C for 2 h. The DNA fragments were purified by ethanol precipitation and analyzed by agarose gel electrophoresis.

### Immunofluorescence

For staining cells on 2D culture, cells were rinsed with PBS once and fixed with 4% paraformaldehyde (PFA) in PBS for 15 min. Cells were washed three times with PBS for 5 min followed by permeabilizing with 0.5% Triton X-100/PBS for 5 min. After rinsing once with PBS, cells were blocked with 1% BSA/0.3% Triton X-100/PBS for 1 h. Next, cells were incubated in primary antibodies diluted in the blocking buffer at 4°C overnight followed by incubating in the secondary antibodies for 1 h R.T. After washing once with PBS, nuclei were counter-stained with DAPI (1:1000 in PBS) for 5 min followed by mounting.

For staining cells on 3D culture, cells were fixed with 2% PFA in PBS supplemented with 50 mM glycine for 30 min at R.T. After washing three times for 20 min with PBS supplemented with 50 mM glycine, cells were blocked with 10% normal goat serum in the IF buffer (0.1% BSA, 0.2% Triton X-100, 0.05% Tween-20 in PBS) at R.T. for 1 h. Primary antibodies were diluted in the blocking buffer and subjected to cells. After overnight incubation at 4°C, cells were washed with the IF buffer for 20 min three times and followed by incubating with secondary antibodies (diluted in the blocking buffer) for 1 h at R.T. After washing once with the IF buffer, nuclei were counter stained with DAPI for 5 min, followed by mounting.

### Immunohistochemistry

Immunostaining of mouse lungs was performed as previously described.(Gao et al., 2016) In brief, tissues were fixed in 10% neutral buffered formalin for 24 hours, embedded in paraffin, sectioned, and stained with hematoxylin and eosin (H&E) for histopathological evaluation. For immunohistochemistry, unstained slides of sectioned tissues were deparaffinized and epitope/antigen retrieval was performed with 10 mM sodium citrate (pH 6.0). The slides were treated with 3% H2O2 to block endogenous peroxidase. The slides were then incubated in 5% normal goat serum/5% BSA/0.3% Triton X-100/PBS and stained with the appropriate primary antibodies. After an overnight incubation in primary antibodies, slides were stained with secondary antibodies and hematoxylin before mounting with the Vectashield mounting medium (Vector Laboratories).

### Western blot

Protein lysates were harvested with RIPA lysis buffer (Sigma Aldrich; R0278) containing protease inhibitor cocktail (Thermo Fisher Scientific; 78429), phosphatase inhibitor 2 and 3 (Sigma Aldrich; P5726and P0044). Samples were standardized for protein concentration using the Pierce BCA protein assay (VWR 23227), and incubated at 100 °C for 10 min under reducing conditions. After denaturation, samples were separated by Bolt 4–12% Bis-Tris Plus Gels (Thermo Fisher Scientific NW04125BOX) and transferred onto a PVDF membrane using iBlot Transfer Stacks (Thermo Fisher Scientific IB401001). Blots were blocked with 5% bovine serum albumin (BSA) in TBST for 1 h at room temperature. Blots were then probed with different primary antibodies (listed in **Table S10**) in the blocking buffer overnight at 4 °C. Blots were washed with TBST for 10 min three times before incubation with secondary antibodies [Anti-rabbit IgG-HRP (Cell Signaling Technology 7074S, 1:5000), Anti-mouse IgG-HRP (Cell Signaling Technology 7076S, 1:5000), anti-goat IgG-HRP (Santa Cruz Biotechnology sc-2354, 1:5000)] in 2.5% BLOT-QuickBlocker (G Biosciences 786-011) in TBST for 1 h at room temperature. Blots were washed with TBST and imaged using chemiluminescent substrate [Pierce ECL (Thermo Fisher Scientific 32209), SuperSignal West Pico PLUS (Thermo Fisher Scientific 34577), or SuperSignal West Femto (Thermo Fisher Scientific 34096)] on the ChemiDox XRS + (Bio-Rad).

### Real time PCR analysis

Total RNA was extracted using the Maxwell® RSC simplyRNA Cells and RNase-free DNase Kit (Promega; AS1390) using an automated RNA/DNA isolation instrument (Maxwell, Promega). cDNAs were synthesized using High-Capacity cDNA Reverse Transcription Kit (Applied Biosystems; 4368814). cDNA corresponding to approximately 10 ng of starting RNA was used for one reaction. qRT-PCR was performed with PowerUp SYBR Green Master Mix (Applied Biosystems) on QuantStudio™ 7 Flex Real-Time PCR System. All quantifications were normalized to endogenous GAPDH. Primer sequences used to amplify genes were purchased from Sigma-Aldrich and are listed in **Table S11**.

### Bulk RNA-sequencing and analysis

Total RNA was isolated as described above. RNA-sequencing was performed by the BGI sequencing facility in Hong Kong. RNA quality, including RNA concentration, RIN value, 28S/18S ratio, and fragment length distribution was confirmed with an Agilent 2100 Bio analyzer (Agilent RNA 6000 Nano Kit). Libraries were prepared by using the standard methodology and run on a BGISEQ-500 platform. Raw reads were quality-checked and subsequently mapped to the mouse genome (mm10_UCSC) through the Spliced Transcripts Alignment to a Reference (STAR)(Dobin et al., 2013), which outputs a BAM file containing both the sequence and its genomic coordinate. Together with a General Transfer Format (GTF) the file containing all the genomic annotations were used to determine the total number of reads per transcript using the HTSeq Count program.(Anders et al., 2015) Differential gene expression was analyzed by using the DESeq2 algorithm(Love et al., 2014) with |fold change| ≥ 2 and FDR < 0.05. Gene set enrichment analysis (GSEA) was performed on a pre-ranked gene list that was generated based on the gene expression changes between sh*Med4* and control cells(Subramanian et al., 2005). Gene expression data were transformed in logarithmic scale and normalized accordingly. Heatmaps showing the gene expression differences between control and *Med4* KD cells were generated by using the Morpheus online tool (https://software.broadinstitute.org/morpheus/).

### ATAC-sequencing and analysis

Chromatin accessibility assays utilizing the bacterial Tn5 transposase were performed as described with minor modifications (Buenrostro et al., 2013). Cells (5.0 x 10^4^) were lysed and incubated with transposition reaction mix for 30 minutes at 37°C. Samples were PCR-amplified and sequenced on an Illumina NextSeq 500. Bulk ATAC-seq raw data were pre-processed using the nfcore/atacseq pipeline version 2.0 (10.5281/zenodo.2634132), implemented in Nextflow (version 22.10.2). Reads were trimmed for both quality and Illumina adapter sequences using ‘trim_galore’ then aligned to mouse assembly mm39 with BWA using the default parameters. Aligned reads with the same start site and orientation were removed using the Picard tool MarkDuplicates (http://broadinstitute.github.io/picard). Density profiles were created by extending each read to the average library fragment size and then computing density using the BEDTools suite (http://bedtools.readthedocs.io). Reads were not extended when generating ATAC-seq read density. Enriched regions were discovered using MACS2 and scored against matched input libraries (|fold change| > 2 and p-value < 0.005). Peaks were then filtered against genomic ‘blacklisted’ regions and filtered peaks within 500 bp were merged to create a full peak atlas. Raw read counts were tabulated over this peak atlas using featureCounts (http://subread.sourceforge.net). All genome browser tracks and read density tables were normalized to a sequencing depth of ten million mapped reads. Peaks were annotated using linear genomic distance, with a gene assigned to a peak if it was within 50 kb upstream or downstream of the gene start or end, respectively. Motif signatures were obtained using the ‘de novo’ approach in Homer v4.7.2 (http://homer.ucsd.edu). The ‘DESeq2’ package in R was employed to identify differential gene expressions and accessibility with the threshold of absolute log_2_ FoldChange > log_2_ (1.5) and FDR ≤ 0.1. When a single gene was linked to multiple ATAC-seq peaks, all associated peaks were retained for analysis. A contingency plot was used to visualize the integrated results from ATAC-Seq and RNA-Seq data. Each quadrant in the plot indicates the statistical significance of either high or low expression for gene expression or accessibility.

### ChIP-seq Analysis

Chromatin immunoprecipitation was performed as described (Terranova et al., 2018) with optimized shearing conditions and minor modifications. The antibodies used were: H3K4me1 (Abcam ab8895), H3K27ac (Abcam ab4729), H3K4me3 (Abcam ab8580), H3K9me3 (Abcam ab3594), H3K27me3 (Abcam ab8898). Sequencing reads were aligned to the mouse reference genome (mm39) using BWA-MEM. SAM files were then converted to BAM format, sorted, and indexed using SAMtools to streamline subsequent analysis. To reduce the impact of PCR duplicates, duplicate reads were marked and removed. Normalized coverage tracks were generated using the bamCoverage command from deepTools, applying RPKM normalization to account for differences in sequencing depth and facilitate direct comparisons between samples. Peak calling was conducted with MACS2 (q < 0.05), employing narrow peak detection for histone marks such as H3K4me3 and H3K27Ac, and broad peak detection for marks like H3K9me3, H3K4me1, and H3K79me2. Input DNA served as a control during peak calling. Heatmaps were generated using deepTools. Chromatin states were identified using ChromHMM to define combinatorial chromatin state patterns based on the histone modifications studied(Ernst and Kellis, 2012). ChIP-seq peaks were annotated using ChIPseeker, and super-enhancers were identified with HOMER based on H3K4me1 and H3K27Ac ChIP-seq signals.

### Statistics and reproducibility

GraphPad 8 software and R were used for statistical analysis. Results were represented as means ± standard deviations (SDs). The normality of the population distribution will be assessed by Shapiro-Wilk test (a significance level of α = 0.05) followed by visualizing with Q-Q plot or histogram. For *in vitro* studies, statistical significance between groups was performed by using an unpaired Student’s t-test with a minimum of three biologically independent samples. For *in vivo* studies, between-group differences were assessed using the Mann-Whitney U test as a nonparametric method with a minimum of five biologically independent samples. Statistical significance was determined when p value was less than 0.05.

## Acknowledgements

This study is dedicated to the memory of our mentor, teacher, and friend, Filippo Giancotti, a pioneer in the metastasis field known for his brilliance, rigor, and kindness. This work was supported by NIH grant R35 CA197566 and partially by CPRIT Recruitment of Established Investigators Award RR160031 (F.G.G.). Use of the Core Facilities was supported by NIH grants P30 CA008748 (MSKCC), P30 CA016672 (MDACC), and P30 CA013696 (Columbia University).

## Author contributions

S.-Y.B. designed the project and performed most of experiments. F.G.G directed the project and provided supervision. S.-Y.B., Y.C., and H.C. performed bioinformatic analyses. D.K. helped in some of the experiments of RT-qPCR, and data analysis. S.-Y.B., A.D.V., R.A.D. were responsible for data analysis and their interpretation. S.-Y.B., A.D.V. and R.A.D. wrote the manuscript.

## Materials & Correspondence

Correspondence and requests for materials should be addressed to Seong-Yeon Bae (S.-Y.B.) and Ronald A. DePinho (R.A.D)

## Extended Data Figure Legends

**Figure S1.**
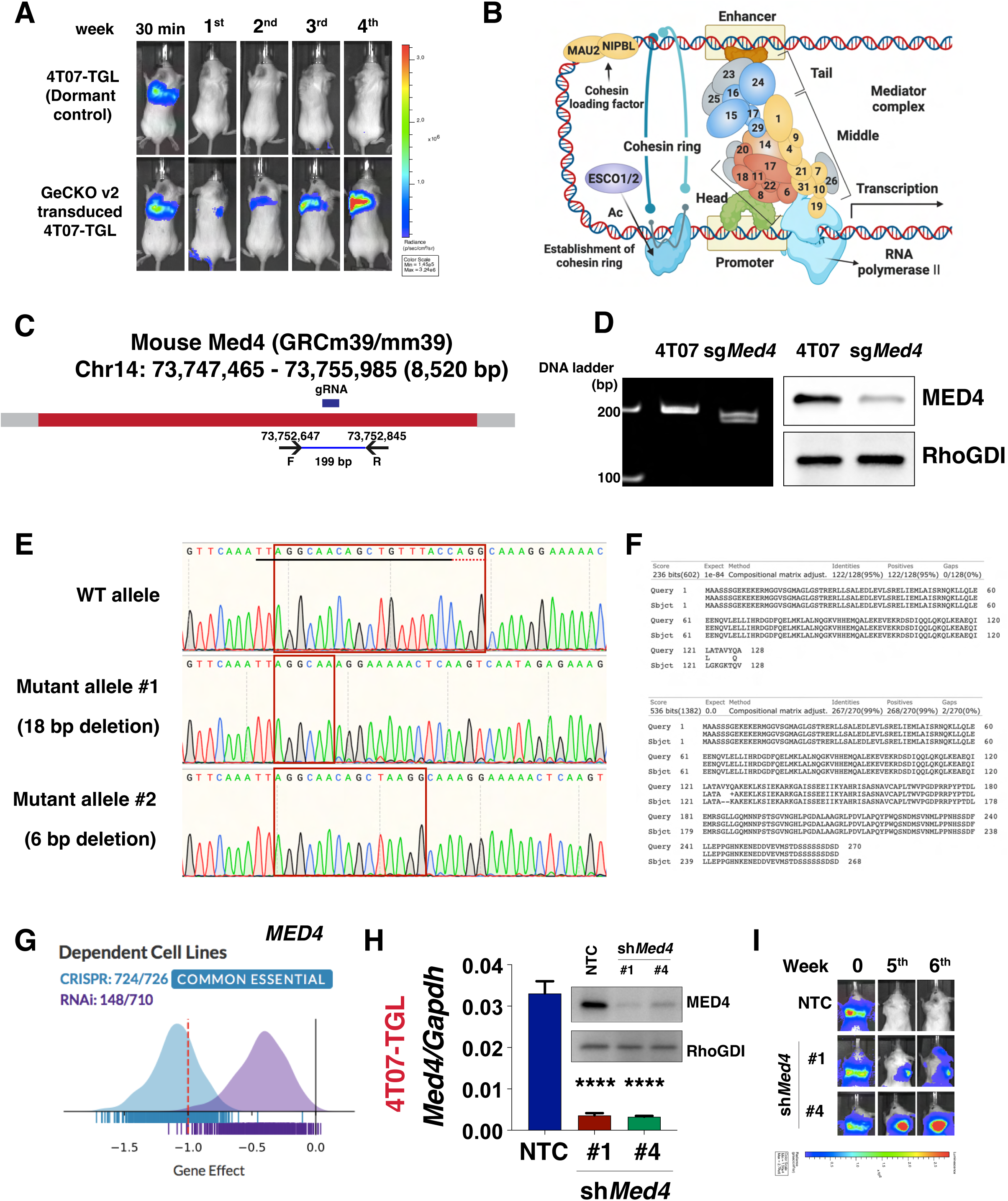
*Med4* was heterozygously deleted to induce metastatic reactivation. **(A)** 4T07-TGL cells either mock control or expressing GeCKO v2 library were inoculated *i.v.* into BALB/c syngeneic mice. Lung metastasis was measured by bioluminescent (BLI) imaging at the indicated time. **(B)** The scheme of chromatin looping for enhancer-promoter interaction. **(C)** Schematics of the mouse chromosomal region encompassing *Med4*, location of the sequence complementary to the gRNA isolated from the screening, and the primer pairs used for genomic DNA PCR. **(D)** The isolated clone from the screening which harbors a gRNA targeting *Med4* was subjected to an indel analysis using genomic DNA PCR, electrophoresed in a DNA-PAGE gel (left), and immunoblotting (right). RhoGDI was served as the loading control. **(E)** Sequences of the fragments of DNA deleted by CRISPR-Cas9. Red boxes indicate the junction. **(F)** Sequence alignment results of translated amino acids from each allele to wild type *Med4*. **(G)** DepMap analysis of the dependency of tumor cell line panels in CRISPR (blue) and RNAi (violet) databases on *MED4*. X-axis: dependency scores. **(H)** RT-qPCR analysis and immunoblots of *Med4* expression in 4T07-TGL cells expressing MirE shRNA targeting either non-targeting control (NTC) or *Med4.* RhoGDI was served as the loading control.; Mean values (± S.D.) p-values; unpaired Student’s t-test. **(I)** 4T07-TGL cells transduced with either NTC or *Med4* shRNAs were inoculated *i.v.* in syngeneic mice. Lung metastasis was measured by bioluminescent (BLI) imaging at the indicated time.

**Figure S2.**
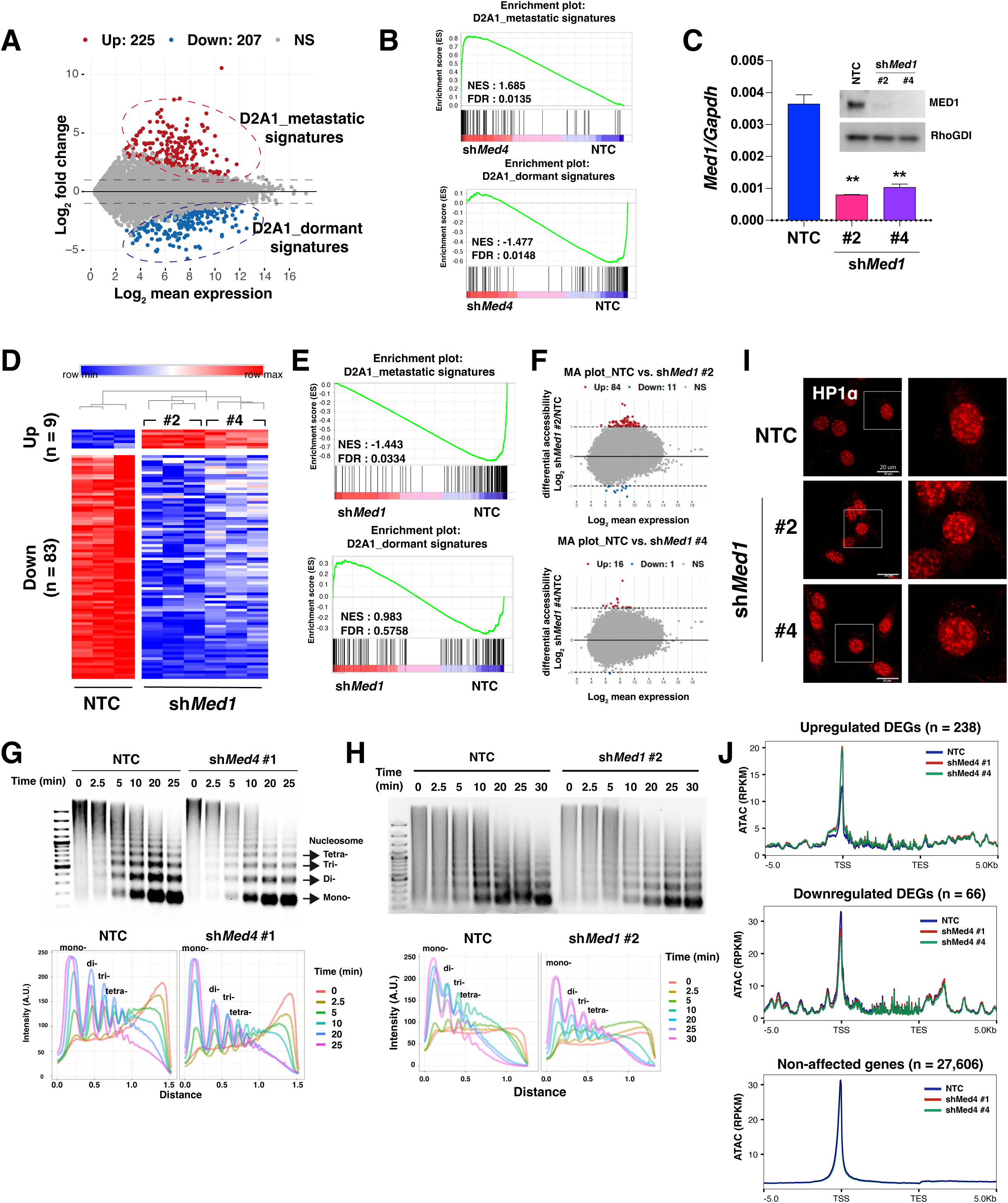
*Med1* alters the transcriptional and epigenetic landscape discordantly with *Med4* while *Med4*-happloinsufficiency driven transcriptional program is necessary for metastasis. **(A)** MA plot from RNA-seq analysis in D2A1 versus D2A1-d cells (Fold change ≥ 2, FDR < 0.05) **(B)** GSEA plots of D2A1_metastatic and D2A1_dormant signatures in Med4-depleted cells as compared with control cells. **(C)** RT-qPCR analysis and immunoblots of *Med1* expression in D2A1-d-TGL cells expressing MirE shRNA targeting either non-targeting control (NTC) or *Med1*; Mean values (± S.D.) p values; unpaired Student’s t-test. **(D)** Heatmap of differentially expressed genes (DEGs; Fold change ≥ 2, FDR < 0.05) in NTC versus *Med1*-silenced D2A1-d-TGL cells. **(E)** GSEA plots of D2A1_metastatic and D2A1_dormant signatures in *Med1*-silenced cells as compared with control cells. **(F)** MA plot of ATAC-seq results comparing NTC versus *Med1*-silenced D2A1-d-TGL cells. Differential accessibility (log_2_ fold change in reads per accessible region) plotted against the mean reads per region. **(G and H)** MNase digestion assays were performed on nuclei isolated from the indicated *Med4*-depleted (G) and *Med1*-depleted (H) cells over increasing incubation times. The panels display representative images (upper) and corresponding quantification graphs (lower). **(I)** D2A1-d cells expressing either NTC or *Med1* shRNAs were stained with HP1α (red). **(J)** ATAC-seq profiles were generated for Up DEGS (up), Down DEGs (middle), and non-affected DEGs (bottom). The profiles display ATAC-seq signal from 5 kb upstream of the TSS to 5 kb downstream of the TES, capturing the chromatin accessibility landscape across gene bodies.

**Figure S3.**
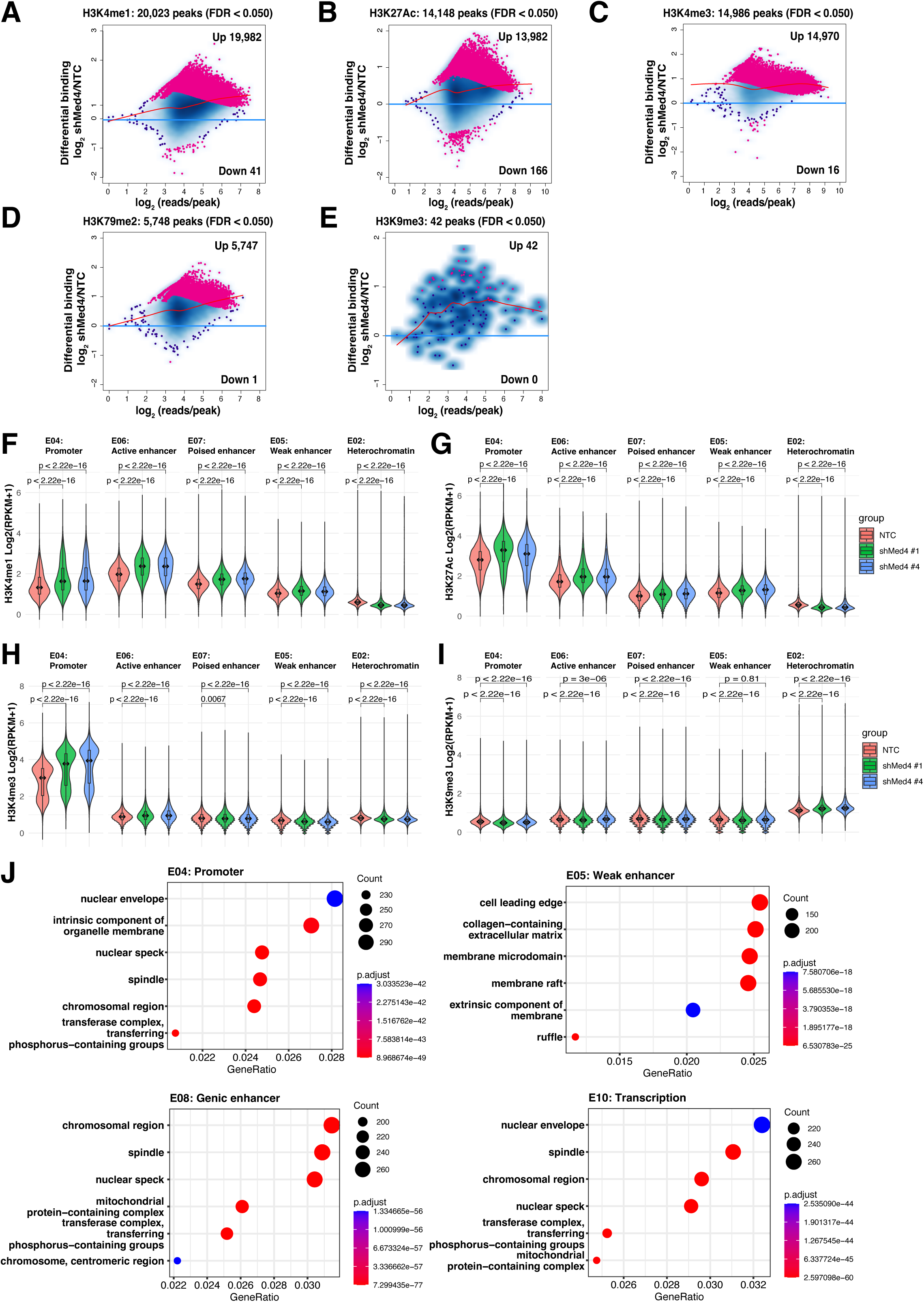
*Med4* haploinsufficiency induces epigenomic reprogramming, making genome permissive to transcriptional activation. **(A-E)** Differential binding sites (FDR < 0.05) were plotted against the mean reads per region for H3K4me1 (A), H3K27Ac (B), H3K4me3 (C), H3K79me2 (D), and H3K9me3 (E). **(F-I)** Violin plots illustrating the distribution of RPKM values for (F), H3K27Ac (G), H3K4me3 (H), and H3K9me3 (I) at each chromatin states defined by ChromHMM. Wilcoxon rank-sum test was used for statistical analysis. **(J)** The top GO cellular component terms in promoter (E04), weak enhancer (E05), genic enhancer (E08), and transcription (E10).

**Figure S4.**
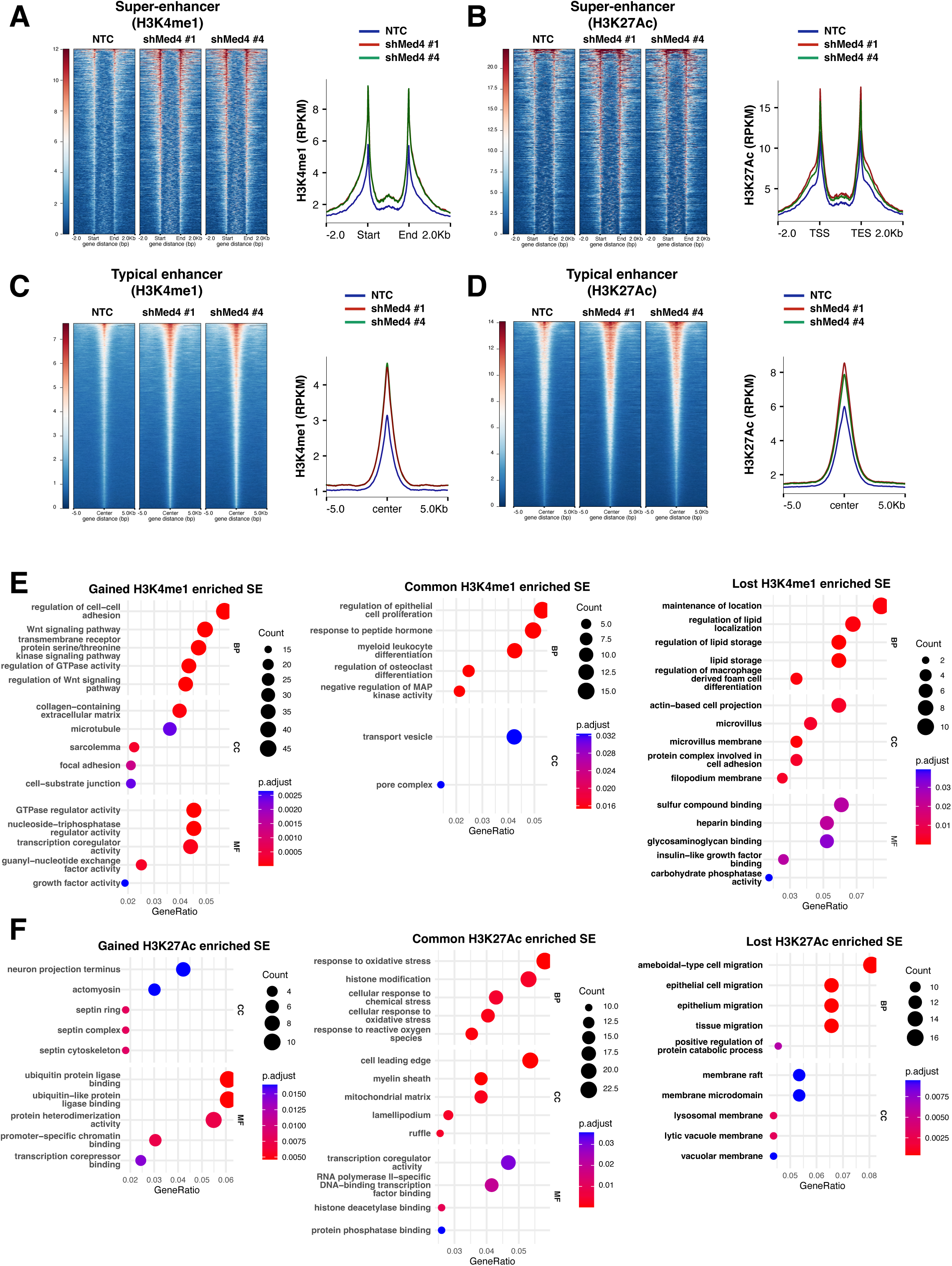
*Med4* reduces epigenomic signals at super-enhancers and typical enhancers genome-wide. **(A** and **B)** Heatmaps (left panels) and average intensity curves (right panels) of ChIP-seq reads (RPKM) for H3K4me1 (A) and H3K27ac (B) at super-enhancer regions, including their flanking 2-kb regions. **(C** and **D)** Heatmaps (left panels) and average intensity curves (right panels) of ChIP-seq reads for H3K4me1 (C) and H3K27ac (D) at typical enhancer regions, displayed within a 10-kb window centered on the middle of the enhancer. **(E and F)** Top 5 GO terms across biological processes (BP), cellular components (CC), and molecular functions (MF) in H3K4me1-enriched super-enhancers (E) and H3K27Ac-enriched super-enhancers (F). Categories are shown for *de novo* super-enhancers in *Med4*-silenced cells (left), persistent super-enhancers shared between *Med4*-silenced and control cells (middle), and lost super-enhancers in *Med4*-silenced cells (right).

**Figure S5.**
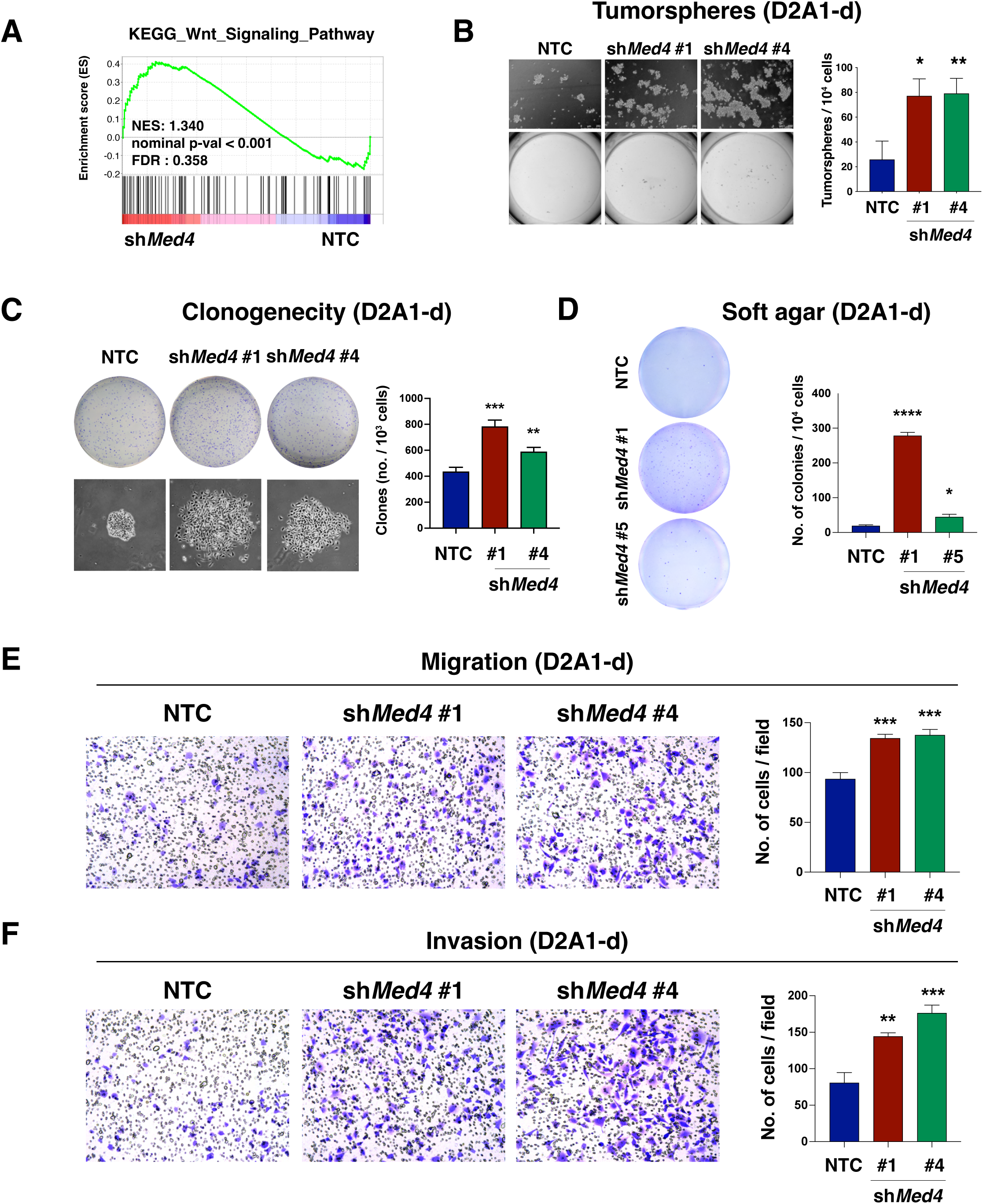
*Med4* suppresses cancer stem cell traits in D2A1-d cells. **(A)** GSEA of KEGG_Wnt signaling pathway signatures. NES, normalized enrichment score; FDR, false discovery rate (n = 3 biological replicates per condition). **(B)** Control or *Med4*-depleted D2A1-d-TGL cells were subjected to tumorsphere assay for 10 days. The panel shows representative images of wells and microscopic images of tumor spheres (left) and mean numbers of organoid (± S.D.; right) per 10,000 cells. p values; unpaired Student’s t-test (n = 3 per group, *P = 0.0122, **P = 0.0089). **(C)** Indicated D2A1-d-TGL cells were subjected to clonogenicity assay for 6 days. The panel shows representative images of dishes and microscopic images of colonies (left) and mean numbers of colonies (± S.D.; right) per 1,000 cells. p values; unpaired Student’s t-test (n = 3 per group, **P = 0.0056, ***P = 0.0006). **(D)** Indicated D2A1-d-TGL cells were subjected to soft agar assay for 10 days. The panel shows representative images of wells and microscopic images of colonies (left) and mean numbers of colonies (± S.D.; right) per 10,000 cells. p values; unpaired Student’s t-test (n = 3 per group, *P = 0.0118, ****P < 0.0001). **(E and F)** Control or *Med4*-silenced D2A1-d-TGL cells were subjected to transwell migration assay (E) and transwell invasion assay (F) for 24 hours. The panel shows representative images of fields (left) and mean numbers of cells (± S.D.; right) per field. p values; unpaired Student’s t-test (E; n = 3 per group, ***P = 0.0007, ***P = 0.0009, F; n = 3 per group, **P = 0.0018, ***P = 0.0007).

**Figure S6.**
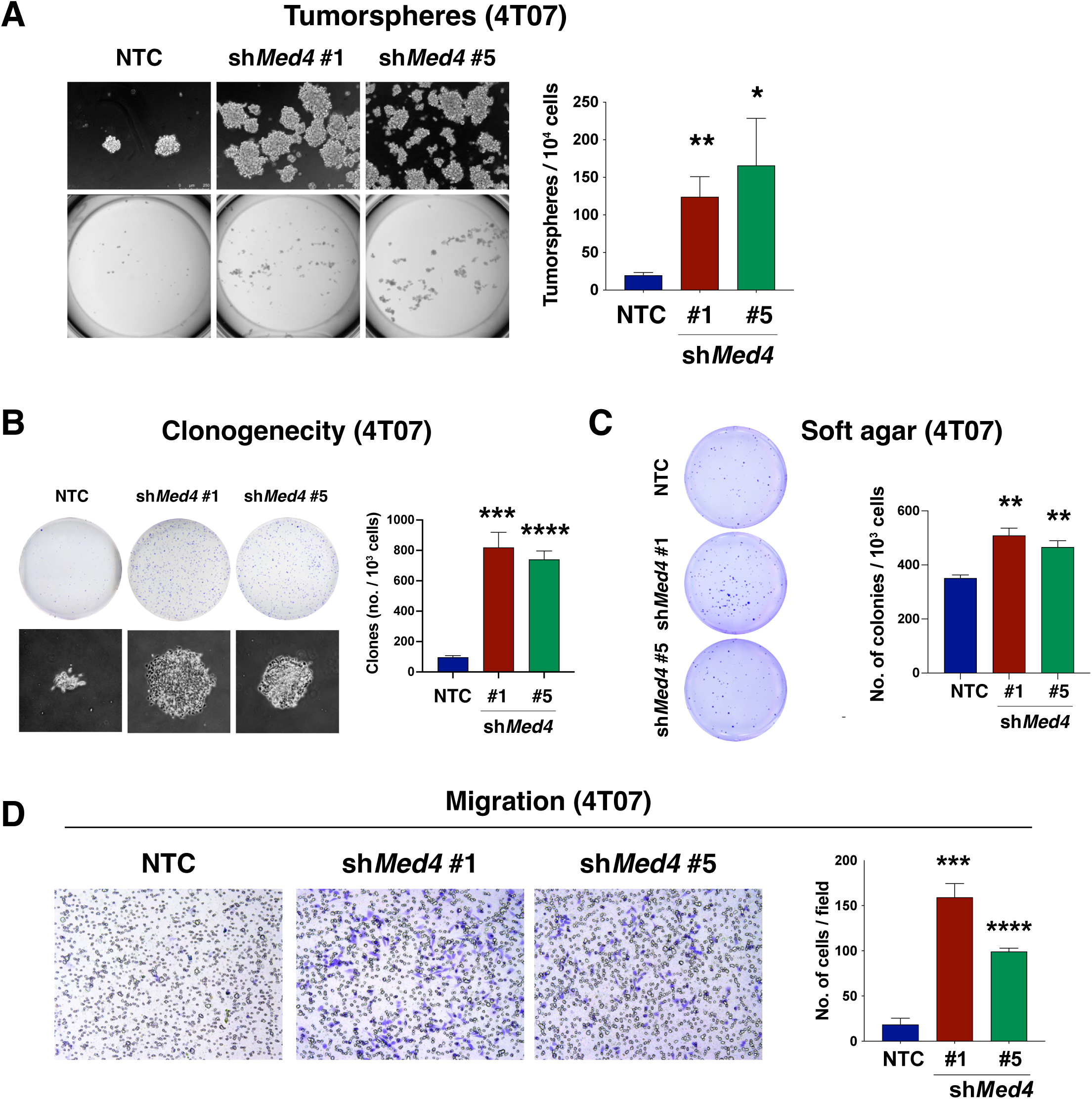
*Med4* inhibits cancer stem cell traits in 4T07 cells. **(A)** Tumorsphere assay results in control and *Med4*-silenced 4T07-TGL cells 10 days post-seeding. The panel shows representative images of wells and microscopic images of tumor spheres (left) and mean numbers of organoid (± S.D.; right) per 10,000 cells. p values; unpaired Student’s t-test (n = 3 per group, *P = 0.0158, **P = 0.0055). **(B)** Indicated 4T07-TGL cells were subjected to clonogenicity assay for 6 days. The panel shows representative images of dishes and microscopic images of colonies (left) and mean numbers of colonies (± S.D.; right) per 1,000 cells. p values; unpaired Student’s t-test (n = 3 per group, ***P = 0.0002, ****P < 0.0001). **(C)** Soft agar assay results in the indicated 4T07-TGL cells 10 days post-seeding. The panel shows representative images of wells and microscopic images of colonies (left) and mean numbers of colonies (± S.D.; right) per 1,000 cells. p values; unpaired Student’s t-test (n = 3 per group, **P = 0.0018, **P = 0.0027). **(D)** Transwell migration results of control and *Med4*-silenced 4T07-TGL cells 24 hours post-seeding. The panel shows representative images of fields (left) and mean numbers of cells (± S.D.; right) per field. p values; unpaired Student’s t-test (n = 3 per group, ***P = 0.0001, ****P < 0.0001).

**Figure S7.**
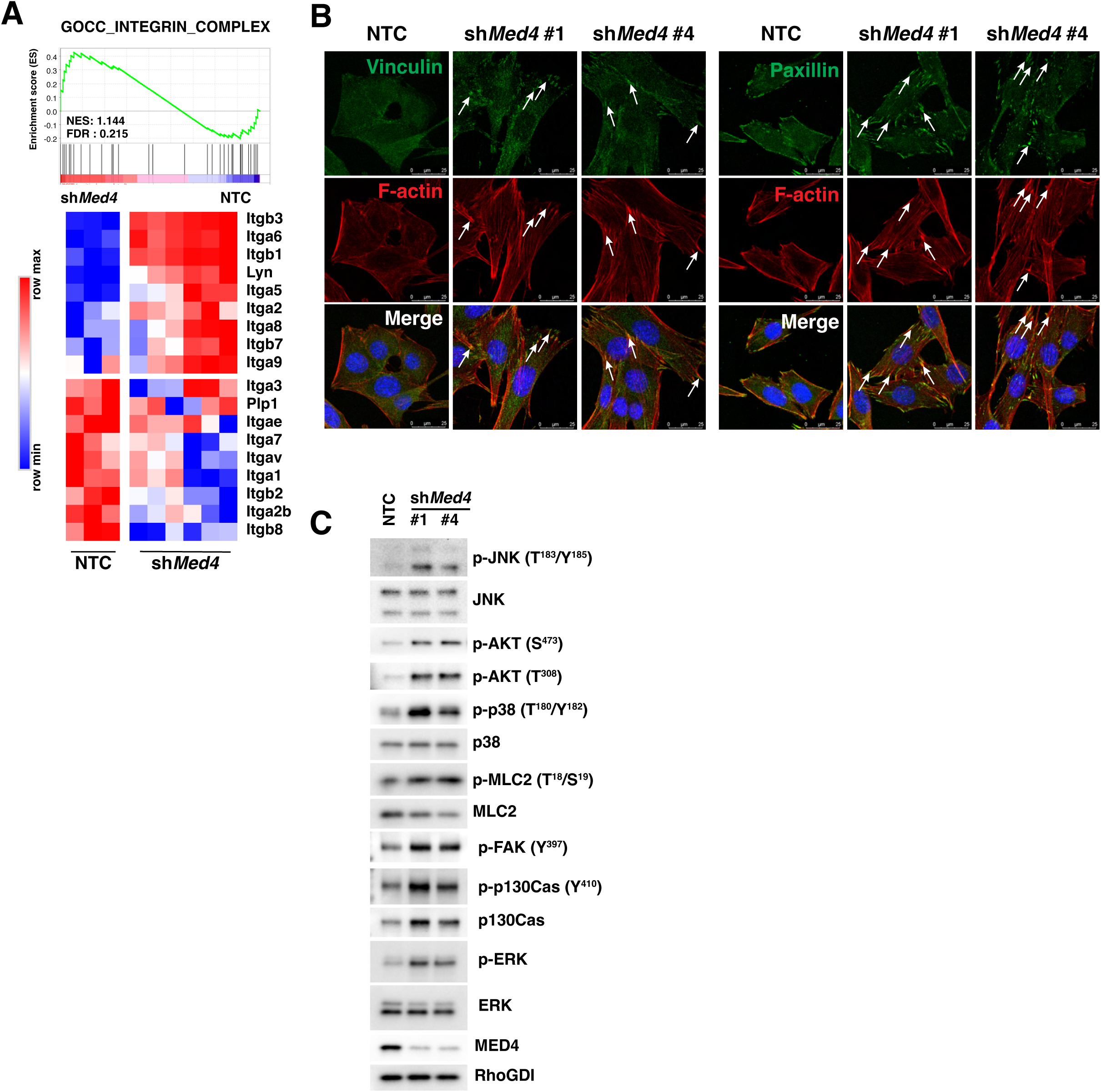
*Med4* alters the repertoire of integrin subunits and mediates the formation of focal adhesion plaques. **(A)** A GSEA plot of integrin complex (GO_CC; top). Heatmap shows the level of expression of integrin subunit genes amongst the top differentially expressed genes in control versus *Med4*-silenced D2A1-d-TGL cells (bottom). **(B)** (Individual channel images for Figure 4C) D2A1-d cells expressing MirE shRNA targeting Med4 cultured on 2D were stained for either vinculin or paxillin (green), F-actin (red), and the nuclei were counterstained with DAPI (blue). Arrows highlight focal adhesion plaques. **(C)** Total cell lysates were subjected to immunoblotting for integrin-signaling molecules. RhoGDI and total ERK, JNK, and p38 were served as internal loading controls.

**Figure S8.**
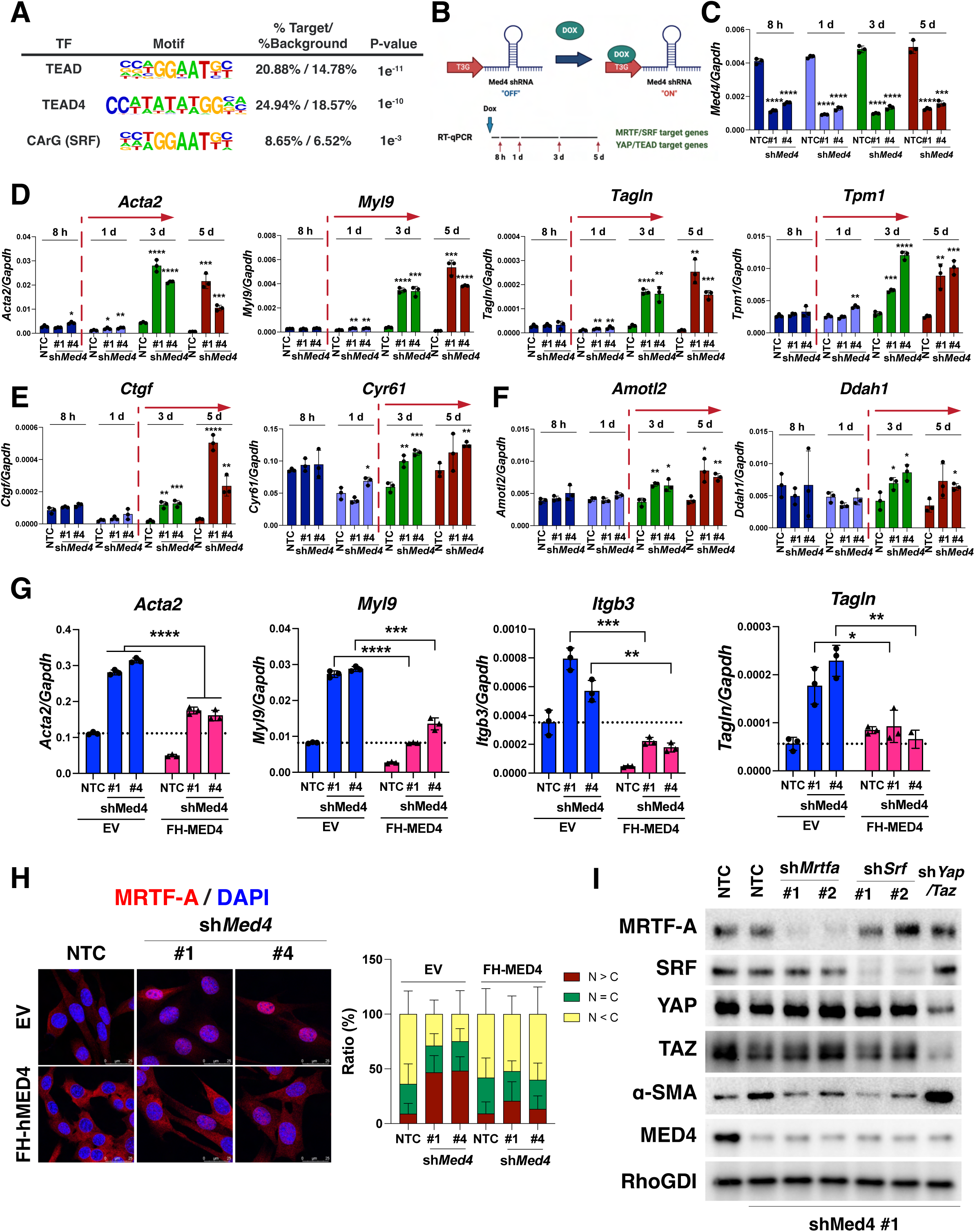
*Med4* mediates the signaling pathway of integrin-mediated mechanotransduction. **(A)** HOMER motif analysis of genes in the upper right quadrant of (Figure 3D) which represents the intersection of RNA-seq and ATAC-seq data in *Med4*-silenced D2A1-d-TGL cells compared to non-targeting control (NTC). p-values were generated using the binomial test. **(B-F)** Schematic representation of inducible expression experiment (B), RT-qPCR results were indicated as relative mean expression of *Med4* (C), MRTF/SRF only target genes (D), shared target genes (E), and YAP/TEAD only target genes (F) and in either control or *Med4*-silenced D2A1-d cells at the indicated time points after initiating doxycycline treatment. **(G)** RT-qPCR results were indicated as relative mean expression of MRTF/SRF only target genes in D2A1-d cells expressing either NTC or Med4 shRNAs were co-transduced with either empty vector (EV) or Flag-HA tagged MED4 cDNA. p values; unpaired Student’s t-test (n = 3 per group, *P = 0.0449, **P = 0.0016, ***P = 0.001, ****P < 0.0001). **(H)** D2A1-d cells expressing either NTC or *Med4* shRNAs were co-transduced with either empty vector (EV) or Flag-HA tagged *MED4* cDNA to rescue MED4 expression. Cells were stained with MRTF-A (red) and counter-stained with DAPI (blue). The panel shows representative images (left) and quantification of subcellular localization (right). N stands for the nucleus and C stands for the cytosol. **(I)** D2A1-d-TGL cells were co-transduced with MirE shRNA targets for *Med4* and MirE shRNA targets the mechanosensitive transcription factors as indicated. Total lysates were subjected to immunoblotting with the indicated antibodies. RhoGDI was served as the loading control.

**Figure S9.**
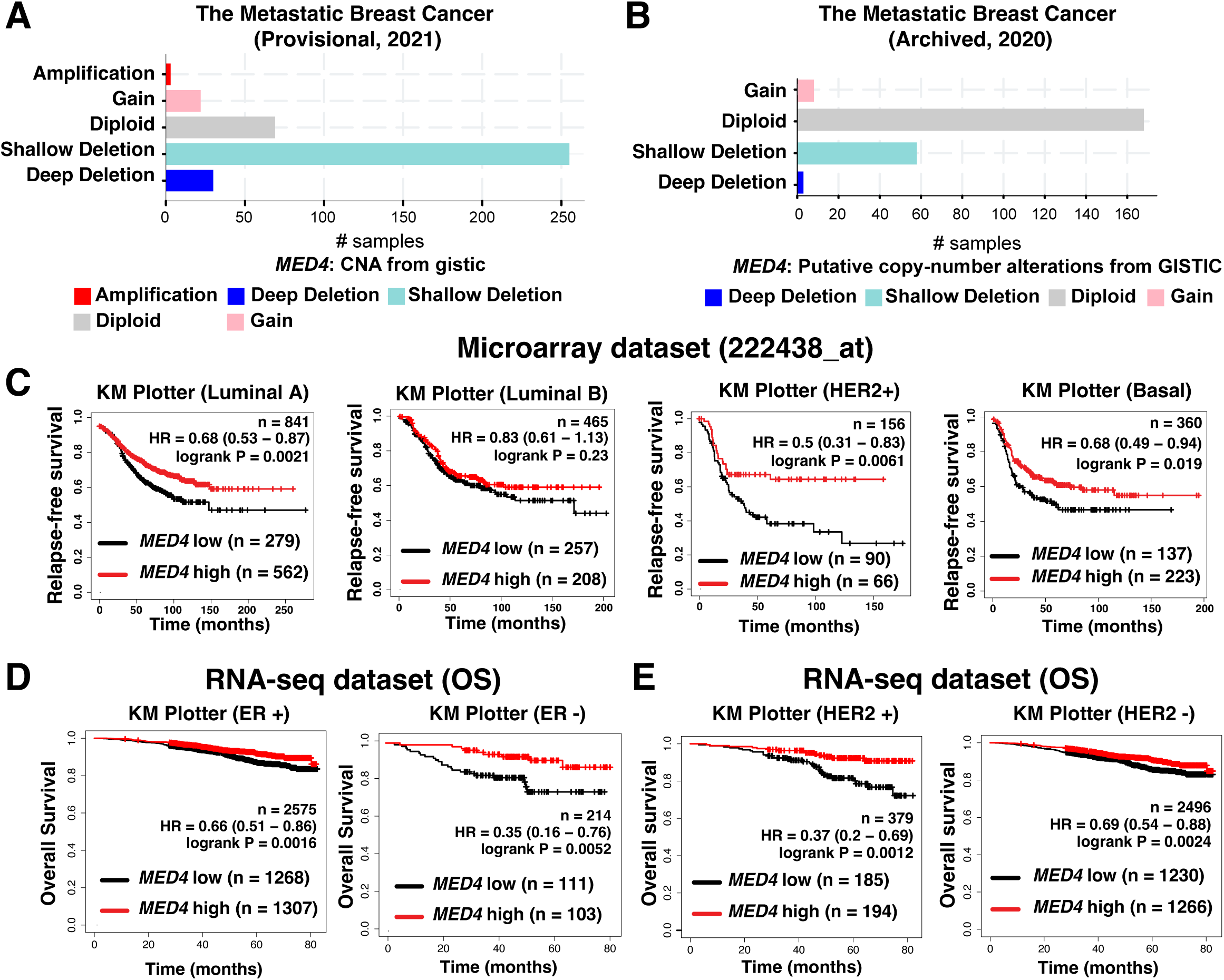
*MED4* serves as a predictor for breast cancer relapse. **(A and B)** cBioportal analysis of copy number alterations in metastatic breast cancer patients from the Metastatic Breast Cancer Project; Provisional (A) and Archived (B). **(C)** Kaplan-Meier analysis of RFS in patients stratified according to *MED4* expression using a probe set, 222438_at as indicated. Patients were subdivided by breast cancer subtypes according to PAM50. **(D** and **E)** KM plotter analysis of overall survival (OS) based on *MED4* expression level from RNA-seq data of breast cancer patients. Patients were subdivided by estrogen receptor (ER) expression level (D) or by human epidermal growth factor receptor 2 (*HER2*) expression level determined by Affymetrix microarray (E). HR, hazard ratio.

